# Machine learning approaches identify immunologic signatures of total and intact HIV DNA during long-term antiretroviral therapy

**DOI:** 10.1101/2023.11.16.567386

**Authors:** Lesia Semenova, Yingfan Wang, Shane Falcinelli, Nancie Archin, Alicia D Cooper-Volkheimer, David M Margolis, Nilu Goonetilleke, David M Murdoch, Cynthia D Rudin, Edward P Browne

## Abstract

Understanding the interplay between the HIV reservoir and the host immune system may yield insights into HIV persistence during antiretroviral therapy (ART) and inform strategies for a cure. Here, we applied machine learning approaches to cross-sectional high-parameter HIV reservoir and immunology data in order to characterize host-reservoir associations and generate new hypotheses about HIV reservoir biology. High-dimensional immunophenotyping, quantification of HIV-specific T cell responses, and measurement of genetically intact and total HIV proviral DNA frequencies were performed on peripheral blood samples from 115 people with HIV (PWH) on long-term ART. Analysis demonstrated that both intact and total proviral DNA frequencies were positively correlated with T cell activation and exhaustion. Years of ART and select bifunctional HIV-specific CD4 T cell responses were negatively correlated with the percentage of intact proviruses. A Leave-One-Covariate-Out (LOCO) inference approach identified specific HIV reservoir and clinical-demographic parameters, such as age and biological sex, that were particularly important in predicting immunophenotypes. Overall, immune parameters were more strongly associated with total HIV proviral frequencies than intact proviral frequencies. Uniquely, however, expression of the IL-7 receptor alpha chain (CD127) on CD4 T cells was more strongly correlated with the intact reservoir. Unsupervised dimension reduction analysis identified two main clusters of PWH with distinct immune and reservoir characteristics. Using reservoir correlates identified in these initial analyses, decision tree methods were employed to visualize relationships among multiple immune and clinical-demographic parameters and the HIV reservoir. Finally, using random splits of our data as training-test sets, machine learning algorithms predicted with approximately 70% accuracy whether a given participant had qualitatively high or low levels of total or intact HIV DNA. The techniques described here may be useful for assessing global patterns within the increasingly high-dimensional data used in HIV reservoir and other studies of complex biology.

## Introduction

The advent of ART has significantly decreased morbidity and mortality associated with HIV infection. However, a long-lived proviral reservoir precludes a cure of infection (1–4). While a number of cell types have been proposed as contributing to the HIV reservoir, the largest and most well-characterized reservoir is within CD4 T cells. Molecular analyses of residual HIV proviruses in CD4 T cells during ART have shown that the reservoir is comprised of a heterogeneous mixture of full-length replication-competent proviruses and a larger set of defective proviruses containing internal deletions or APOBEC-mediated hypermutation(s) (5–7). Infected cells can undergo clonal expansion and contraction over time in response to homeostatic cues or antigen-driven activation (8–10). In addition to the presence of a latent HIV reservoir, PWH on therapy exhibit a persistent level of T cell immune activation and other altered phenotypes compared to uninfected individuals, involving cells of both the innate and adaptive immune systems (11). Specifically, HIV infection is associated with increased levels of inflammatory biomarkers and increased immune cell activation (12–15). PWH also exhibit a depleted naïve T cell compartment (16) and increased expression of activation/exhaustion and senescence markers such as CD38, PD-1 and KLRG1(17–21). The molecular mechanisms underlying persistent immune activation and dysfunction for PWH on ART are unclear, but it is possible that ongoing viral gene expression and reverse transcription generates ligands for pattern recognition receptors in the innate immune system (22). However, another important potential cause of ongoing immune activation during ART is persistent sequelae of the massive disruptions to T cell homeostasis, intestinal permeability and lymphoid structures that occur during untreated HIV infection (23–25).

It is possible that the HIV reservoir and the host immune system interact in multiple complex ways during therapy. The dynamics of how the HIV reservoir affects the immune system and vice versa are important to characterize and may lead to novel ideas to promote reservoir depletion or to inhibit immune dysfunction that may be related to ongoing HIV expression in persistently infected cells. A number of previous studies have examined the relationship between reservoir size and the immune system (21,26–44). Consistent major findings from these studies include enrichment of the HIV reservoir in CD4 T cells expressing activation and exhaustion markers, positive correlation of the reservoir with central memory T cells (Tcm) and transitional memory (Ttm) T cell subsets, and inverse correlation of the HIV reservoir with the CD4 nadir and the CD4/CD8 T cell ratio. Recent studies indicate enrichment of the intact HIV reservoir in cells expressing VLA-4, but no correlation with classic T cell activation markers (26, 36). One important limitation of most of these studies is that they were not able to distinguish intact proviruses from defective proviruses, making the relationship between the immune system and the intact reservoir unclear. Furthermore, most of these studies examined a limited number of surface markers and may have missed associations that would be revealed by a higher-resolution examination of immunophenotypes. Additional assessment of immune correlates of the intact HIV reservoir in additional cohorts is thus needed. Finally, there is a recent, growing appreciation for the profound influence of ART initiation timing and length of treatment on the size and composition of the HIV reservoir that must be accounted for in assessing persistent immune activation in PWH on therapy (11, 45).

In this study, we use a mixture of traditional statistical approaches and novel data science methods to assess the relationship between the immune system and the HIV reservoir across a cross-sectional study of 115 PWH. We identify numerous significant correlations between immune cell populations and the size of the total HIV reservoir, as well as smaller number of associations with the intact reservoir, including CD127 expression in CD4 T cells. This study highlights the potential of machine learning approaches to visualize global patterns in high-parameter studies of the HIV reservoir. Additionally, these approaches corroborate recent findings in the HIV persistence field regarding preferential intact proviral decay and immune dynamics and highlight the complex immunologic signatures of HIV latency.

## Results

### Cohort description, reservoir quantification, and immunophenotyping

This cross-sectional study evaluated 115 participants with HIV infection for at least one year and who had been receiving ART for at least 0.9 years (median 9 years, IQR 5.2-16.6) (**Table 1**). The study cohort was 77 percent male, 60 percent black and median participant age was 45 years.

**Table 1:**
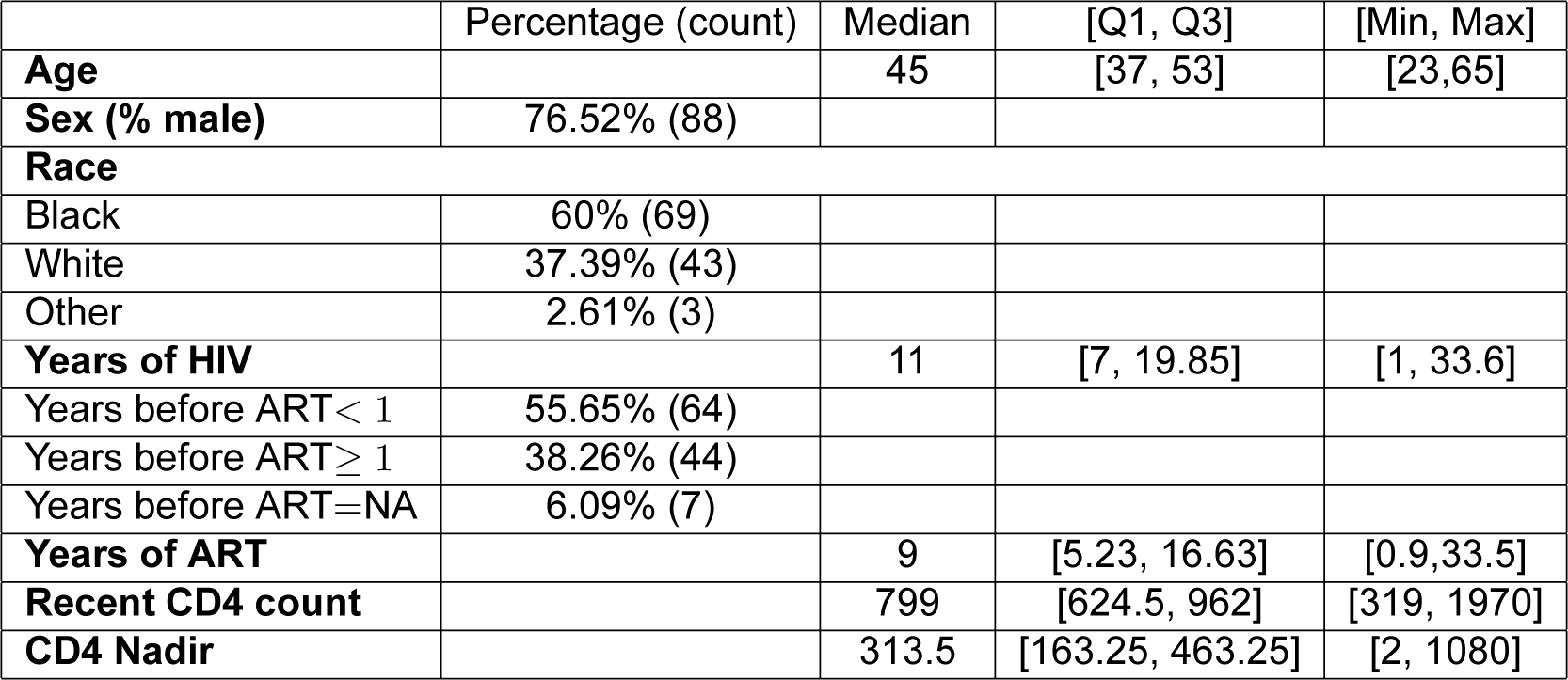
Participant demographic and clinical characteristics. For demographics and clinical information, we report percentage for categorical variables, medians and [Q1, Q3] for real-value variables. ART is antiretroviral therapy. CD4 counts reported in cells/mm^3^. Years of HIV has 1 missing value, Years of ART has 7, and CD4 Nadir has 3; consequently, these missing values are not included in median and quantiles computations. Years before ART means years of HIV infection before ART initiation.

|  | Percentage (count) | Median | [Q1, Q3] | [Min, Max] |
| --- | --- | --- | --- | --- |
| **Age** |  | 45 | [37, 53] | [23,65] |
| **Sex (% male)** | 76.52% (88) |  |  |  |
| **Race** |  |  |  |  |
| Black | 60% (69) |  |  |  |
| White | 37.39% (43) |  |  |  |
| Other | 2.61% (3) |  |  |  |
| **Years of HIV** |  | 11 | [7, 19.85] | [1, 33.6] |
| Years before ART < 1 | 55.65% (64) |  |  |  |
| Years before ART ≥ 1 | 38.26% (44) |  |  |  |
| Years before ART=NA | 6.09% (7) |  |  |  |
| **Years of ART** |  | 9 | [5.23, 16.63] | [0.9,33.5] |
| **Recent CD4 count** |  | 799 | [624.5, 962] | [319, 1970] |
| **CD4 Nadir** |  | 313.5 | [163.25, 463.25] | [2, 1080] |

The Intact Proviral DNA assay (IPDA) was performed on isolated total CD4 T cells (46). The IPDA estimates of the frequency of total proviruses, intact proviruses, and the percent intact of total proviruses for each participant. Immunophenotyping was performed using 25-color spectral flow cytometry on peripheral blood mononuclear cells (PBMCs) from each study participant. A representative flow gating is shown in **Figure S1**. Briefly, cells within the live lymphocytes gate were first defined as T cells (CD3+/CD56-), NK cells (CD3-/CD56+), or NKT cells (CD3+/CD56+). T cells were then further subdivided as CD4+ or CD8+ T cells, and then as naïve T cells (Tn, CD45RA+/CCR7+), central memory T cells (Tcm, CD45RA-/CCR7+), effector memory cells (Tem, CD45RA-/CCR7-) and terminally differentiated effector cells (Teff, CD45RA+/CCR7-). Expression of various surface markers within each of these subsets was then examined (CD38, HLA-DR, PD-1, KLRG1, CD127, CD27, NKG2A). In addition to measuring the baseline abundance of surface proteins, we also stimulated PBMCs from each participant with or without a pool of HIV-derived peptides and measured intracellular cytokines TNF*α*, IL-2, and IFN*γ* as well as the surface expression of the degranulation marker CD107a. HIV-specific T cell responses were defined as the difference in frequency of cytokine/CD107a+ cells between HIV stimulated and nonstimuated cells. Frequencies of CD4 or CD8 T cells that were single- or double-positive for TNF*α*, IL-2, IFN*γ*, or CD107a were used for analysis.

These analyses resulted in the determination of a total of 144 parameters for the cohort of n = 115 PWH on ART. The 144 parameters include 133 immunophenotypic cell frequencies, 3 HIV reservoir parameters (frequency of intact HIV DNA, frequency of total HIV DNA, and percentage intact of total HIV DNA), and 8 clinical-demographic parameters. Clinical-demographic variables (**Table 1**) included age, biological sex, years of ART, estimated years of HIV infection prior to ART (here, we use categorical variables: years before ART=NA, years before ART*<* 1, years before ART*≥* 1), CD4 nadir, most recent CD4 T cell count, and race (Caucasian, African-American, other). No significant differences in the reservoir based on biological sex or race were observed (**Table S1**). The median and interquartile range (IQR) for all immune cells population variables are presented in **Table S2**.

### Correlation of total, intact, and percent intact HIV DNA with immunophenotypes

For the initial analysis, Spearman correlation of the clinical-demographic and immunophenotypic data with HIV reservoir metrics was assessed. 69 host variables correlated (unadjusted p *<* 0.05) with one or more characteristics of the HIV reservoir (**Table S3**). Following Benjamini-Hochberg p-value adjustment for multiple comparisons using a false discovery rate correction at 5%, 31 variables were found to be correlated with total HIV DNA, 3 with intact HIV DNA, and 4 with the percentage of intact HIV DNA (**Table 2**). Of note, the correlations observed were generally weak to moderate in magnitude (the mean of the absolute values of Spearman correlation coefficients was 0.31 for total HIV DNA, 0.33 for intact HIV DNA, and 0.35 for percent intact HIV DNA).

**Table 2:** PWH features correlate with HIV reservoir characteristics. The abundance of 144 immune cell populations were determined by flow cytometry and the HIV reservoir was quantified by intact proviral DNA assay for a cohort of 115 people with HIV (PWH). Each abundance and clinical and demographic variable was correlated with total HIV reservoir frequency, intact reservoir frequency and the percentage of intact proviruses. Spearman correlation coefficients (**bold**) are shown for 36 variables that had significant p-values (*<* 0.05) after Benjamini–Hochberg correction for multiple comparisons. Each feature/subset is ranked by the absolute value of the correlation coefficient for the total reservoir frequency. For years of ART, we compute correlation based on 108 participants, excluding participants with missing years of ART values.

| Variable | Total | Intact | Percent intact |
| --- | --- | --- | --- |
| %CD8 T | **0.4052** | **0.3562** | 0.0068 |
| %CD38+/HLA-DR- CD4 T | **-0.3891** | -0.1098 | 0.2664 |
| %KLRG1-/PD-1- CD4 T | **-0.3808** | -0.1289 | 0.2334 |
| %Tn CD4 T | **-0.3802** | -0.2031 | 0.1714 |
| %NKG2A+ CD4 T | **0.3618** | 0.2904 | 0.0179 |
| %PD-1-/CCR7+ CD4 T | **-0.3590** | -0.1082 | 0.2283 |
| %CD4 T | **-0.3564** | **-0.3195** | -0.0079 |
| %Tcm CD8 T | **0.3466** | 0.1752 | -0.1814 |
| %CD38+ CD4 T | **-0.3366** | -0.0611 | 0.2829 |
| %PD-1-/CCR7- CD4 T | **0.3300** | 0.1824 | -0.0938 |
| %CD38+/HLA-DR- CD8 T | **-0.3267** | -0.0837 | 0.2636 |
| %PD-1+ CD4 T | **0.3222** | 0.0664 | -0.2470 |
| Age | **0.3172** | 0.1669 | -0.1471 |
| CD4 Nadir | **-0.3164** | -0.1512 | 0.1927 |
| %PD-1+ Tn CD4 T | **0.3119** | 0.1246 | -0.0962 |
| %Tn CD8 T | **-0.3028** | -0.2697 | 0.0154 |
| Years of ART | **0.3062** | -0.0688 | **-0.4523** |
| %PD-1+/CCR7+ CD8 T | **0.2926** | 0.1361 | -0.1254 |
| %CD38-/HLA-DR+ CD4 T | **0.2849** | 0.0920 | -0.1500 |
| %KLRG1-/PD-1- CD8 T | **-0.2757** | -0.2182 | 0.0162 |
| %PD-1+ Tn CD8 T | **0.2738** | 0.1411 | -0.0705 |
| %CD38+/HLA-DR- Tn CD8 T | **-0.2676** | -0.0610 | 0.2119 |
| %PD-1+/CCR7+ CD4 T | **0.2665** | 0.0859 | -0.1673 |
| %KLRG1+/CD27+ CD8 T | **0.2606** | 0.2212 | 0.0319 |
| %CD27+ CD4 T | **-0.2602** | -0.0531 | 0.1767 |
| %KLRG1-/CD27+ CD4 T | **-0.2575** | -0.0464 | 0.2171 |
| %HLA-DR+ CD4 T | **0.2565** | 0.1110 | -0.0970 |
| %PD-1+ CD8 T | **0.2541** | 0.1969 | 0.0035 |
| %CD38+ CD8 T | **-0.2418** | 0.0109 | **0.3114** |
| %CD38-/HLA-DR+ Tn CD8 T | **0.2404** | 0.0738 | -0.1346 |
| %Tem CD4 T | **0.2402** | 0.0888 | -0.1669 |
| %CD127+ CD4 T | -0.1298 | **-0.3160** | -0.2539 |
| %CD107a-IFNγ-IL-2+TNFα+ CD4 T | 0.0568 | -0.1775 | **-0.3223** |
| %CD107a-IFNγ+IL-2+TNFα- CD4 T | 0.0285 | -0.2455 | **-0.3265** |

With regard to clinical-demographic variables, CD4 nadir prior to ART initiation was negatively correlated with total HIV DNA (Spearman *r* = −0.32), whereas age (*r* = 0.31) and years of ART (*r* = 0.31) were positively correlated with total HIV DNA (**Table 2**). Years of ART (*r* = −0.45) was negatively correlated with the percentage of intact proviruses (**Table 2**), consistent with the hypothesis that intact proviruses are progressively eliminated from the reservoir over time. We then examined the association between the reservoir and time on ART further (**Figure 1**). Correlation plots of total reservoir frequency, intact reservoir frequency, and percentage intact HIV DNA versus years of ART were visualized (**Figure 1A-C**). The plots showed no significant correlation for intact HIV DNA frequency versus years of ART (**Figure 1B**), while total HIV DNA reservoir frequency was positively correlated with the length of ART treatment. (**Figure 1A**, Spearman *r* = 0.31). When we examined a plot of percent intact proviruses versus time on therapy (**Figure 1C**), we observed a biphasic decay pattern, consistent with previous reports (47–49). Indeed, when we fitted piece-wise linear functions with allowances for up to 2 breaks, the best-fit model had decay slope changes at 1.6 and 9 years of ART. The R^2^ score of a piece-wise linear model with two breaks was 44.18%, while the score for a linear model without any breaks was 16.3%. In PWH within approximately 0-6 years of ART initiation, a significant proportion of the participants exhibited high fractions of intact proviruses (50-90%). After approximately 6 years of ART, however, the majority of participants had reservoirs for which intact proviruses represented a minor fraction of the overall reservoir (0-30%). Of note, several immune features also demonstrated statistically significant correlations with years of ART, including CD8 T cell expression of CD38 (negative correlation), CD8 Tcm frequency (positive correlation), and CD107a+IL-2+IFN*γ*-TNF*α*-CD8 T cells (positive correlation)(**Figures S2, S3, S4, Table S4)**.

**Figure 1:**
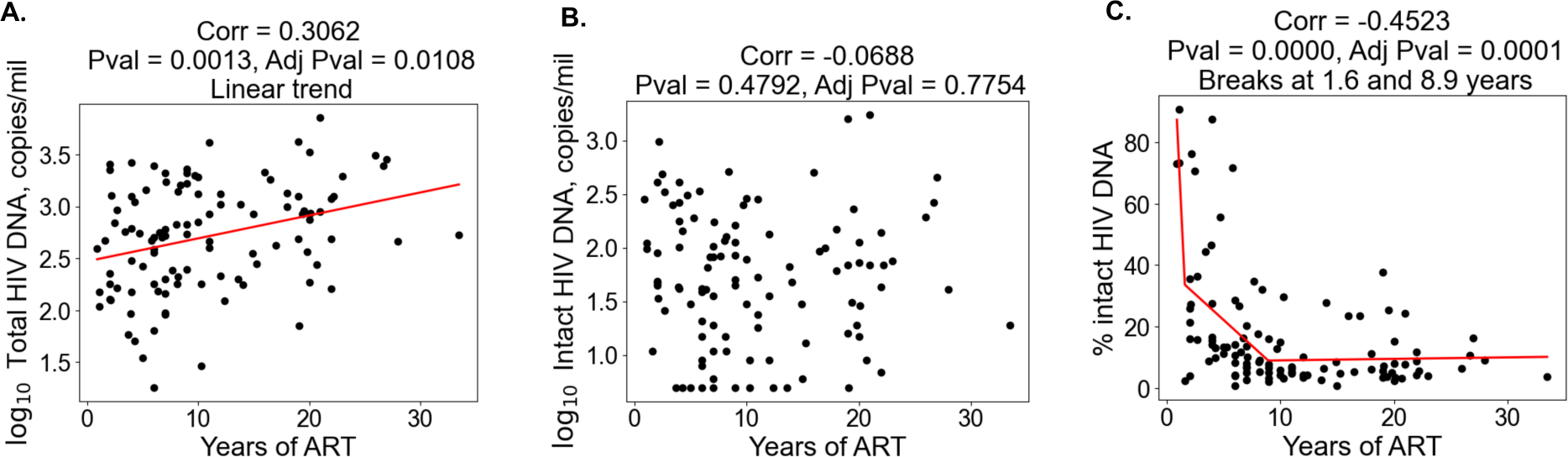
Duration of treatment and the HIV reservoir. Scatterplots for years of ART vs total HIV reservoir frequency **(A)**, intact reservoir frequency **(B)** and percent intact **(C)** are shown. Each dot represents an individual study participant. Correlation coefficients and corresponding p-values are shown for each plot. Participants that have missing values of years of ART were not included in the plot. For percent intact, piece-wise linear function with two breaks is fitted. For total HIV reservoir frequency a linear function is fitted.

Examination of immunophenotypic parameter correlations with HIV DNA metrics demonstrated several notable correlations (**Table 2, Figures S2, S3, S4**). As previously reported, CD4 T cell frequencies were negatively correlated with total HIV DNA frequencies (*r* = −0.36), whereas CD8 T cells were positively correlated (*r* = 0.41). This relationship held true for intact HIV DNA frequencies but not for the percentage of intact HIV DNA (**Table 2**). Frequencies of CD4 and CD8 T naïve cells, as well as their expression of CD38, were significantly negatively correlated with total HIV DNA frequency, consistent with the hypothesis that larger reservoirs are associated with depletion of the naive T cell subsets and increased immune activation. By contrast, CD4 and CD8 Tcm and Tem cells were weakly positively correlated with total HIV DNA, possibly due to increased differentiation of the naive compartment into these memory subsets in people with larger reservoirs. Markers of T cell activation and exhaustion were weakly but significantly positively correlated with total HIV DNA, including NKG2A+ CD4 T, PD-1+ CD4 T, HLA-DR+ CD4 T, PD-1+ CD8 T, and PD-1+/CCR7+ CD8 T frequencies, among others (**Table 2, Figures S2, S3, S4)**. For both intact and total HIV DNA, an inverse correlation with the CD4/CD8 T cell ratio was observed (**Figure S5**). Interestingly, for the percentage of intact HIV DNA, negative correlations were observed with the frequencies of 2 subsets of bifunctional HIV peptide-stimulated CD4 T cells (IL-2+TNF*α*+ and IFN*γ*+IL-2+), as well as with years on ART. Notably, the magnitude of correlation with immune parameters was typically stronger for total HIV DNA compared to intact HIV DNA (**Table 2**). This pattern may reflect a larger influence of total HIV DNA on immunophenotype of the host, due to the overall higher abundance of total HIV DNA relative to the intact reservoir. A notable exception to this pattern was CD127-CD4 T cell frequencies, which were significantly positively correlated with intact but not total HIV DNA.

Overall these findings support the notion that larger HIV reservoirs are associated with increased levels of immune activation and depletion of the naive T cell compartment. Furthermore, the data indicate that there may be some immune features, such as CD127 expression on CD4 T cells that are uniquely associated with the intact reservoir rather than the total reservoir.

### Leave-One-Covariate-Out (LOCO) inference identifies specific HIV reservoir and clinical-demographic parameters important for the prediction of immunophenotypes

Given the evidence that reservoir and immune recovery dynamics likely occur in concert, we next used a variable importance approach - Leave-One-Covariate-Out (LOCO) inference analysis, to account for potential confounding variables and more carefully assess the relative importance of a given variable to an immunophenotype. The LOCO inference approach is described in **Figure 2A**. First, a least-squares regression linear model that predicts the dependent variable is fitted on a set of independent variables, and an R^2^ value is generated (**Figure 2A Step 1**). Next, one independent variable is excluded from the model, the model is refitted, and an adjusted R^2^ value is determined (**Figure 2A Step 2**). Finally, the change in R^2^ scores, Δ*R*^2^, is calculated (**Figure 2A Step 3**). This process is then repeated for all independent variables in the model, and the Δ*R*^2^ values are plotted in a heatmap for the variables of interest (**Figure 2B-D**).

**Figure 2:**
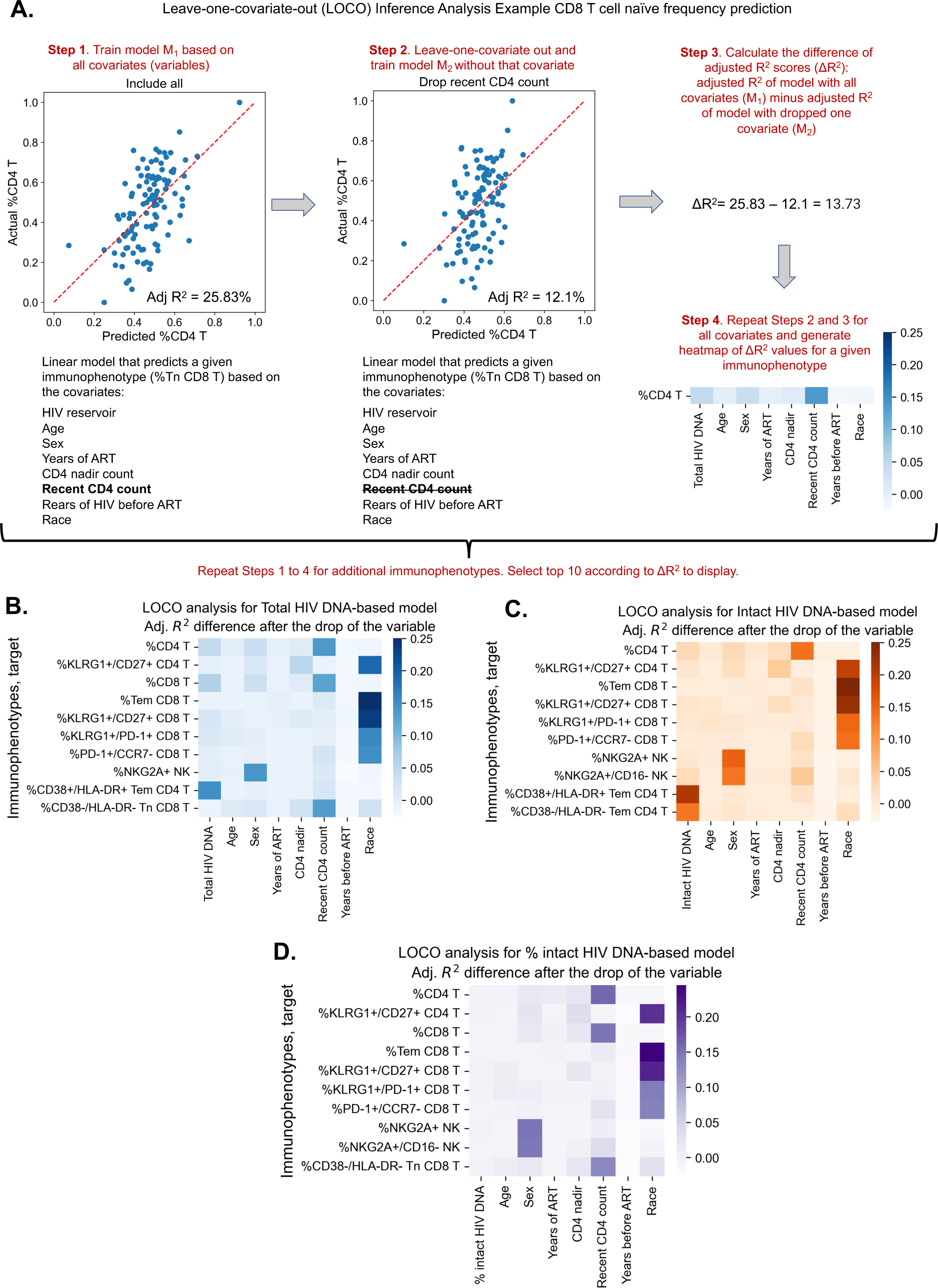
Leave-one-covariate-out (LOCO) analysis for clinical-demographic features and reservoir characteristics while predicting immunophenotypes. **A.** Explanation of leave-one-covariate-out (LOCO) analysis based on example of %CD4 T for clinicaldemographic features and reservoir characteristics while predicting immunophenotypes. Analysis was performed for all 133 immunophenotypes considered in the study. The top 10 biggest drops in adjusted R^2^ scores are reported for models that use total reservoir frequency (**B.**), intact reservoir frequency (**C.**), or percent intact (**D.**) as features in addition to clinical and demographic information. Participants with missing years of ART values are excluded from this analysis. The missing value of the CD4 nadir for one participant is imputed. In **Figure S6**, the LOCO analysis results are visualized for all 133 immunophe-

We applied the LOCO inference analysis to all 133 immunophenotype parameters (dependent variables). Three separate analyses were conducted for a set of independent variables consisting of age, sex, years of ART, CD4 nadir, recent CD4 account, years of HIV infection prior to ART (=NA, *<* 1, *≥* 1), and one of the three HIV reservoir metrics (total HIV DNA, intact HIV DNA, or percentage intact HIV DNA). Multicollinearity analysis confirmed that independent variables from this analysis were not correlated with each other to a degree that would interfere with model generation (**Table S5**). Each analysis produced a heatmap of 133 dependent immunophenotype variables, with visualization of the variables for each immunophenotype (**Figure S6, Tables S6-S8**). For each of these analyses, we displayed the top 10 variables with the largest Δ*R*^2^ value for LOCO inference incorporating either total (**Figure 2B**), intact (**Figure 2C**), or percentage intact (**Figure 2D**) HIV DNA.

The least-squares linear regression models were generally weakly explanatory (*R*^2^ values in range [-11.11%, 35.36%]) of the global variability in immunophenotypes (**Tables S6-S8**, **Figure 2B-D**, **Figure S7**). Across all 3 analyses, age and race emerged as an important covariate of several immunophenotypes related to T cell subsets, particularly the naïve and central memory compartments, as well as KLRG-1, a marker of immune senescence (**Figure 2 and Figure S6**). Interestingly, biological sex was a key variable for model prediction of NK cell NKG2A expression, and race was important for Tem CD8, KLRG1+/CD27+ CD8 frequency for all three models. Recent CD4 count was important for predicting the frequencies of HLA-DR-CD8 Tn cells and CD4, CD8 T cell frequencies, but more so for the models incorporating total HIV DNA and percent intact than the intact HIV DNA frequency (**Figure 2A-C**). Total HIV DNA frequency was an important model input for prediction of PD-1+ CD4 T naïve cells, HIV-specific IL-2+TNF*α*-IFN*γ*-CD107-CD4 T cells, and CD4/CD8 T cell frequencies, among others (**Figure 2B, Figure S6**). Total HIV DNA frequency and intact HIV DNA frequency were also important predictors of CD4 T effector memory HLA-DR+CD38+ expression (**Figure 2B and 2C**).

In general, total and intact HIV DNA were predictive of similar immune features (**Figure S6, Tables S6-S8)**; however, total HIV DNA generally typically had greater magnitude Δ*R*^2^ than intact HIV DNA. The percentage of intact HIV DNA did not appear to contribute greatly to the predictive power of the model for the variance in the immunophenotypes examined (**Figure S6C),** most likely due to its relatively high correlation with years of ART Overall these findings further demonstrate the existence of a number of clinical parameters that are associated with immune signatures in PWH, including age, gender, and years of ART. Additionally, these results highlight key potential confounding variables that should be considered when interpreting the association of virological parameters with the immune system of PWH.

### Receiver operator characteristic curve analysis demonstrates distinct immune parameters associated with intact and total reservoir frequencies

The complex relationship between the HIV reservoir and the immune system led us to evaluate if machine learning (ML) approaches that combine several parameters into models of the HIV reservoir could be a useful approach to understand this interaction. Before this could be attempted, we first identified a defined set of the most valuable variables from which ML models could be built. To achieve this, we first binarized the three HIV reservoir metrics (total HIV DNA frequency, intact HIV DNA frequency, %intact HIV DNA) into high (above median) or low (below median) reservoir groups and generated receiver operator (ROC) curves for the 144 clinical-demographic and immunophenotype parameters (**Figure 3**). The area under the curve (AUC) of the ROC curve indicates the ability of the feature (immune parameter or clinical/demographic information) to correctly identifying a participant as having qualitatively low or high total HIV DNA frequency (**Figure 3A**), intact HIV DNA frequency (**Figure 3B**), or percentage intact HIV DNA (**Figure 3C**). A model that randomly guesses high or low reservoir has an AUC of 0.5. In this analysis, 57 host variables had an AUC higher than 0.6 for one or more of total HIV DNA, intact HIV DNA, or percent intact DNA (**Table S9**). The most effective individual immune markers for classification of high versus low total HIV DNA frequency included %NKG2A+ CD4 T (AUC = 0.70), %PD-1+ Tn CD4 T (AUC = 0.68), and %CD38+/HLA-DR-CD8 T (AUC = 0.68). In contrast, the most effective markers for classifying based on the frequency of intact HIV DNA frequency included %CD127+ CD4 T (AUC = 0.71), %CD8 T (AUC = 0.66), and %Tn CD8 T (AUC = 0.65). Finally, for percentage intact HIV DNA, years of ART (AUC = 0.72), %CD107-IFN*γ*-IL-2+TNF*α*+ CD4 T (AUC = 0.65) and %KLRG1+/PD1+ CD4 T (AUC = 0.65) were the most effective (**Figure 3**). Similar to the findings in the Spearman correlations (**Table 2**) and LOCO inference analysis (**Figure 2**), there were more variables that had AUC values *>* 0.6 for total reservoir frequency (**Table S9**): 44 variables above this threshold for the total reservoir, but only 19 variables for the intact reservoir and 23 variables for percent intact. Overall, this approach allowed us to derive a ranked list of the most predictive immune parameters for each aspect of the HIV reservoir, and these highly ranked features were thus used for subsequent dimension reduction and ML modeling.

**Figure 3:**
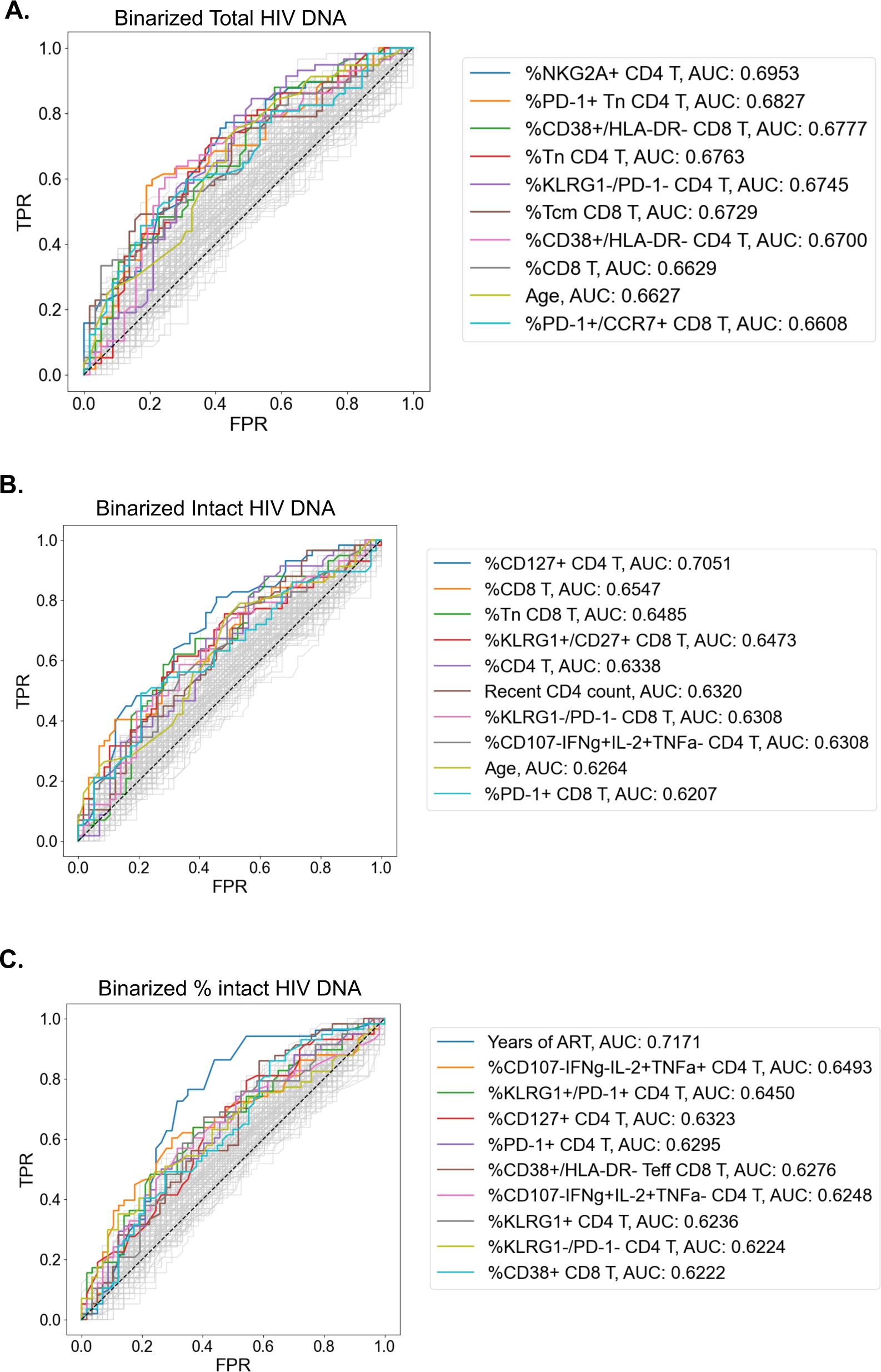
Receiver operating characteristic (ROC) curves identify PWH parameters that can classify reservoir characteristics. For total reservoir frequency (**A**), intact reservoir frequency (**B**), and percent intact (**C**), ROC curves are plotted for all 144 immune markers, demographics, and clinical variables (shown in gray). Axes represent the true positive rate (TPR) and the false positive rate (FPR) for each variable for classifying study participants into low (below median) versus high (above median) reservoir frequency. ROC curves for ten variables with the highest AUC values are shown in color for each HIV reservoir characteristic. Striped black lines represent the ROC curves of a random model. For years of ART ROC curves, we exclude participants with missing years of ART val_2_u_7_es.

### Dimension reduction reveals two clusters of PWH with distinct HIV reservoirs

To examine the overall structure of dataset we employed an unsupervised dimension reduction (DR) machine learning approach (PaCMAP (50)) using the 10 immune cell features with the highest AUC values for classifying participants based on total HIV DNA frequency, intact HIV DNA frequency, or percentage intact HIV DNA (**Figure 3, Table S9**). Interestingly, while no clear clustering was observed when we use intact reservoir frequency or percentage intact HIV DNA associated features, we observed two distinct clusters of PWH (cluster 1 and cluster 2, **Figure 4A**) when using total HIV DNA associated features. These clusters were of roughly equal size with 50 participants in cluster 1 and 65 in cluster 2. Projecting total reservoir frequency (above or below the median) onto the clustering plot, we observe a strong distinction between the two clusters in terms of total HIV DNA reservoir characteristics, (**Figure 4B,C**). Analysis of quantitative reservoir frequencies across clusters (**Figure 4C-E**) demonstrated that cluster 1 is characterized by a smaller total HIV reservoir frequency, but greater percentage of intact proviruses (**Figure 4C-E**). In contrast, cluster 2 is defined by a larger total reservoir frequency, but lower percentage of intact proviruses (**Figure 4C-E**). The frequency of intact proviruses also tended to be lower for cluster 1 but this difference was not significant. We also visualized the dataset using principal component analysis (PCA) (**Figure S9**), and observed that participants from each of the clusters identified by PaCMAP occupied different areas of the plot, although the separation was not as clear as with PaCMAP.

**Figure 4:**
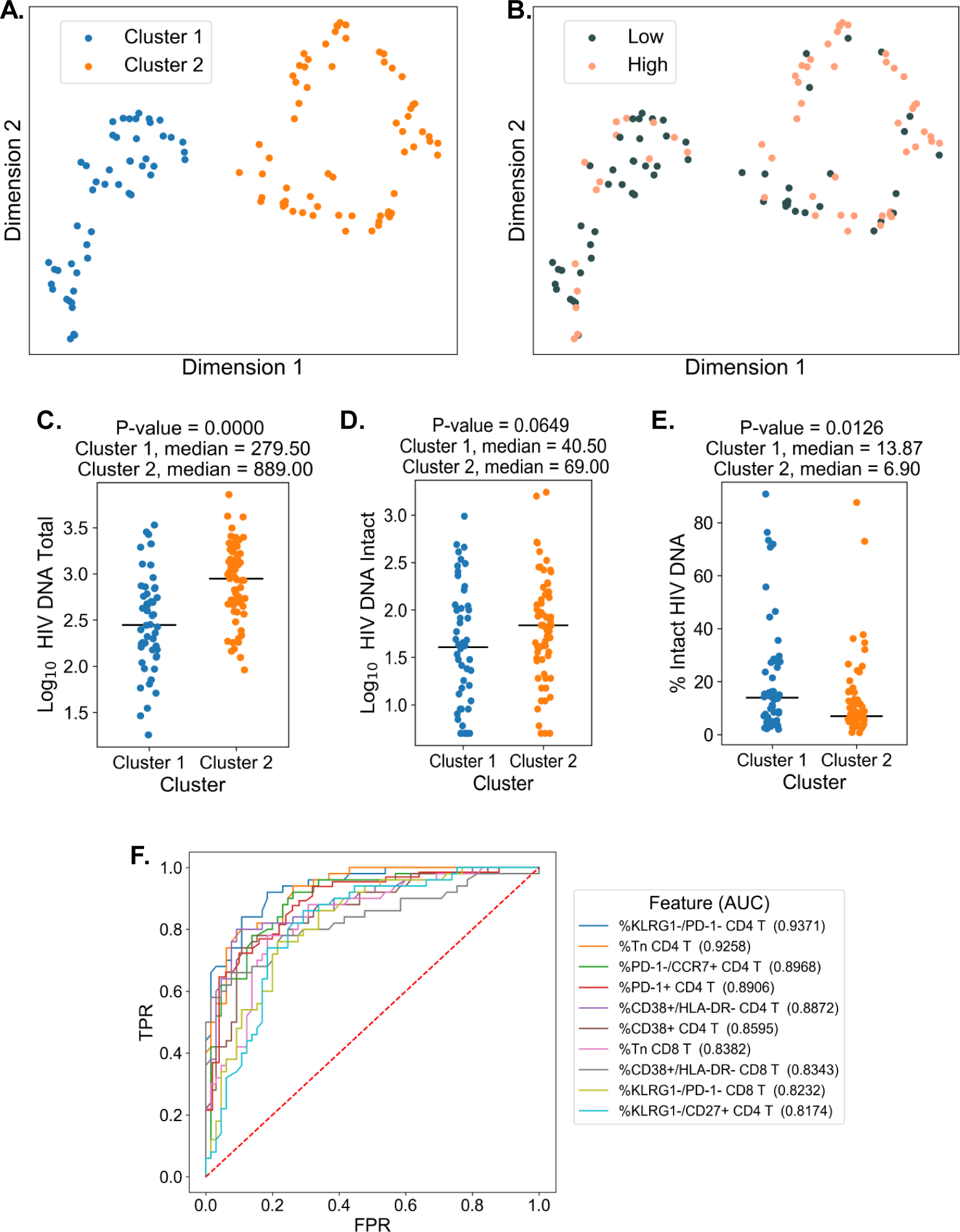
Dimension reduction reveals two major clusters of PWH with distinct immune systems and reservoirs. **A.** PaCMAP was applied to the data using the ten immune cell features with the highest AUC values for classifying participants based on total reservoir frequency, and two clusters (cluster 1 and cluster 2) are identified. **B**. Same as A but data points are color-coded by total reservoir frequency (high=pink, low=gray). Total reservoir frequency (**C.**), intact reservoir frequency (**D.**), and percent intact (**E.**) is shown for participants within each cluster. **F.** Key immune cell features that distinguish cluster 1 from cluster 2 are identified by visualizing the features with the highest AUC values with respect to classifying cohort participants based on cluster membership. Axes represent the true positive rate (TPR) and the false positive rate (FPR) for each variable. Immune markers and clinical-demographic features are shown for each cluster in Figure S8.

To gain insight into the immune cell features that distinguish these two clusters, we generated ROC curves for the 133 immune parameters based on their ability to identify membership in cluster 1 versus cluster 2 for each participant (**Figure 4F**). Many features that distinguished the clusters overlapped with the features selected for the clustering analysis, including expression patterns of CD38, HLA-DR, KLRG-1 and PD-1 on CD4 and CD8 T cells and CD4 T naïve cell frequencies. However, some novel features emerged, including CD4 KLRG1-CD27+ frequency, CD8 KLRG1-PD1-frequency, and CD8 Tn frequency (**Figure 4F and Figure S8**). Cluster 2, which had a higher total and intact absolute reservoir frequency but lower percentage of intact proviruses, was associated with a lower fraction of naïve CD4 and CD8 T cells. Additionally, cluster 2 had generally higher expression of immune exhaustion markers, including KLRG1 and PD-1 (**Figure S8**). These data indicate that cluster membership is highly associated with the overall level of immune activation and exhaustion for the individual. Notably, with regard to other clinical variables, cluster 2 had an older median age, longer time on ART, lower CD4 nadir, and lower current CD4 T cell count (**Figure S8**). We have previous shown that cannabis use is associated with lower levels of immune activation markers in PWH, despite having minimal impact on the size of the HIV reservoir (51). Interestingly, a subset of participants in this study had clinical data regarding cannabis (CB) use. CB users were enriched in cluster 1, which had lower levels of immune exhaustion markers (**Figure S8**). Overall, these data demonstrate that, within the cohort of PWH on long-term ART, there were two major clusters of participants with distinct reservoir characteristics and immunophenotypes.

### Decision tree visualization of PWH with respect to reservoir characteristics

Since the interaction of the immune system and the HIV reservoir is multifactorial, we hypothesized that models that consider multiple parameters simultaneously could more accurately describe the overall dataset, and provide insights regarding the biology of the HIV reservoir and the host immune system. To accomplish this, we employed a decision tree approach to visualize combinations of variables that classify participants as having high (above median) or low (below median) reservoir frequency. Compared to the dimension reduction technique, decision tree visualization is an interpretable supervised approach and does not require post-hoc analysis.

We first selected 35 variables with the highest ROC AUC values for either total reservoir frequency, intact reservoir frequency, or percentage intact HIV DNA to be considered for model generation (**Figure 4**). Using these parameters, we fitted Generalized and Scalable Optimal Sparse Decision Trees (GOSDT) (52) to the data in order to classify high versus low total HIV DNA, intact HIV DNA, or percentage intact HIV DNA. We required the trees to achieve at least 80% accuracy for classifying PWH in the cohort, as well as have at least five PWH in each leaf. Since these trees are based on the entire dataset, these models are thus descriptive rather than predictive.

For the total HIV DNA frequency decision tree, only four immune variables were required to accurately describe high versus low total HIV DNA status (**Figure 5A, Figure S10**): CD8 T cell frequency, CD4 nadir, %CD38+HLA-DR-CD8 T cells, and %NKG2A+ CD4 T cells. The tree divided the cohort into five subgroups (leaves), among which three have high total reservoir frequency and two have low total reservoir frequency. Comparing the labels provided by GOSDT model with the actual data, the tree achieved 83.5% accuracy (i.e. misclassifying 19 PWH among the overall cohort of 115 PWH). Notably, when we combined all samples from “high total reservoir” leaves and all samples from “low total reservoir” leaves, we observed a significant difference in the actual median total reservoir frequencies for these two groups (266/M for low total and 1288.5/M for high total, Mann Whitney U test p-value is 3.56e-13, **Figure 5B**).

**Figure 5:**
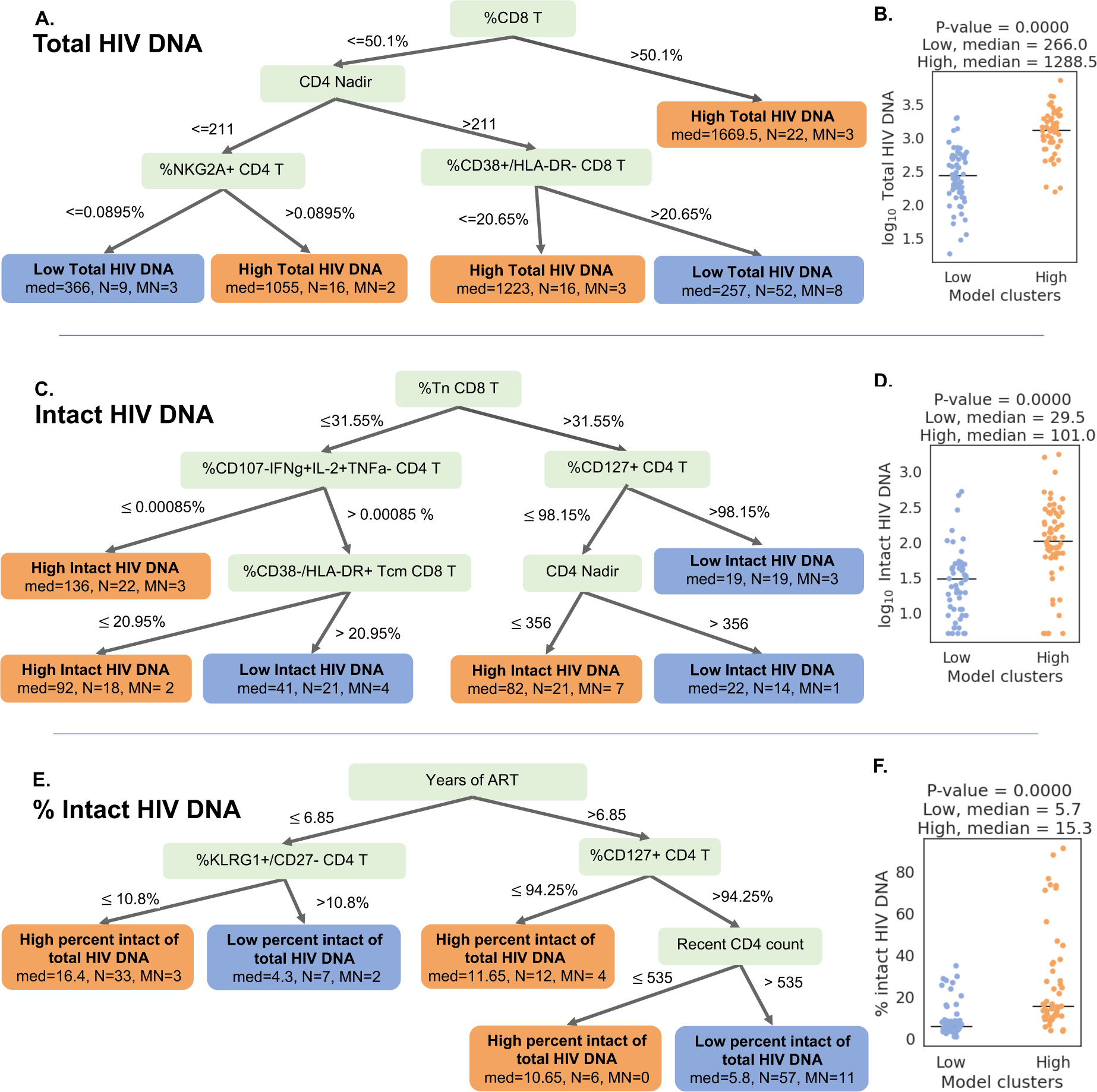
Decision tree visualization of the association of immune cell subsets with reservoir characteristics. **(A, C, E)** Host variables (immune cell frequencies, demographic, and clinical information) were used to visualize the PWH dataset using the optimal sparse decision trees algorithm GOSDT. The overall set of PWH was classified as likely having high (above median, orange “leaves”) or low (below median, blue “leaves”) total reservoir frequency **(A)**, intact reservoir frequency **(C)**, and percent intact **(E)**. In each leaf, “med” denotes the median HIV characteristic of PWH, N is the number of PWH in the leaf, and MN is the number of misslabeled PWH. **(B, D, F)** PWH in model leaves associated with high (orange) or low (blue) reservoir frequency characteristics were aggregated and a Mann-Whitney U test was performed to determine statistical significance between the actual total reservoir frequency of the “high” and “low” groups for total reservoir frequency **(B)**, intact reservoir frequency **(D)**, and percent intact **(F)**. For the percent intact tree we exclude participants with missing values of years of ART. For total and intact reservoir frequency, missing values of years of ART were imputed, however, since the trees do not use this variable, imputations do not influence results. Visualization trees are explained with sets of rules in Supplemental Materials: **Figure S10** for total reservoir frequency, **Figure S11** for intact reservoir frequency, and **Figure S12** for percent intact.

We then repeated this approach for classifying the cohort participants with respect to intact reservoir frequency (**Figure 5C**, **Figure S11**). Despite the overall lower correlation between the immune cell phenotypes and the intact reservoir size, this tree nevertheless achieved 82.6% accuracy. The tree had six leaves (three with low intact reservoir, three with high intact reservoir) and relied on %Tn CD8 T cells, %CD107a-IFN*γ*+IL-2+TNF*α*-CD4 T cells, %CD127+ CD4 T cells, %CD38-HLA-DR+ Tcm CD8 T cells, and CD4 Nadir. When we combined the ‘high intact’ leaves together and the ‘low intact’ leaves together, we observed a significant difference in median intact reservoir frequency between the groups (29.5/M for low intact and 101/M for high intact, Mann Whitney U test p-value is 9.59e-08, **Figure 5D**).

Finally, a third iteration of GOSDT generation was performed, classifying the cohort participants with respect to percentage intact of total proviruses (**Figure 5E**, **Figure S12**). This tree achieved 82.4% accuracy and relied on years of ART treatment, %KLRG1+CD27-CD4 T cells, %CD127+ CD4 T cells, and recent clinical CD4 count. This tree has two leaves with low percent intact reservoir and three leaves with high percent intact reservoir. We observed a significant difference in the actual percent intact values between the model generated high percent intact and low percent intact leaves (median 5.7% for low percent intact leaves and median 15.3% for high percent intact leaves: Mann Whitney U test, p-value is 5.5e-10, **Figure 5F**).

Collectively, these decision trees highlight combinations of immune parameters that can accurately describe qualitatively high versus low total, intact, and percentage intact HIV DNA in a cohort of n = 115 PWH. This visualization serves as a basis for mechanistic hypotheses about the interactions of the immune system and HIV reservoir during long-term ART.

### Machine learning algorithms identify immune feature combinations that predict high versus low reservoir metrics with **∼**70% test accuracy

Although clustering and decision tree analysis permit visualization and understanding of global structures within a dataset, we were curious if combinations of immune and clinical-demographic parameters could actually accurately predict, rather than only visualize, the size of the HIV reservoir. We considered five machine learning algorithms including Logistic Regression with L2 regularization (LR), CART, Support Vector Machines with RBF kernel (SVM), Random Forest (RF), and Gradient Boosted Trees (GBT). Initially, we attempted to predict reservoir frequency as a continuous variable, but found that models trained for this purpose performed poorly and tended to overfit to training sets (**Figure S13**). Thus, we focused our efforts on predicting high (above median) versus low (below median) reservoir frequency. For reservoir characteristics, we measured accuracy of the models over 10 random splits of our data into training and test sets (**Figure 6A and Table S10)**. Overall, we found that for total reservoir frequency, Logistic Regression achieved highest mean classification accuracy (69.31%) in test data. The accuracy of these models is likely limited by the sample size (n=115) and noise in the data, as evidenced by lower *R*^2^ score of machine learning models for non-binarized HIV characteristics (**Figure S13**).

**Figure 6:**
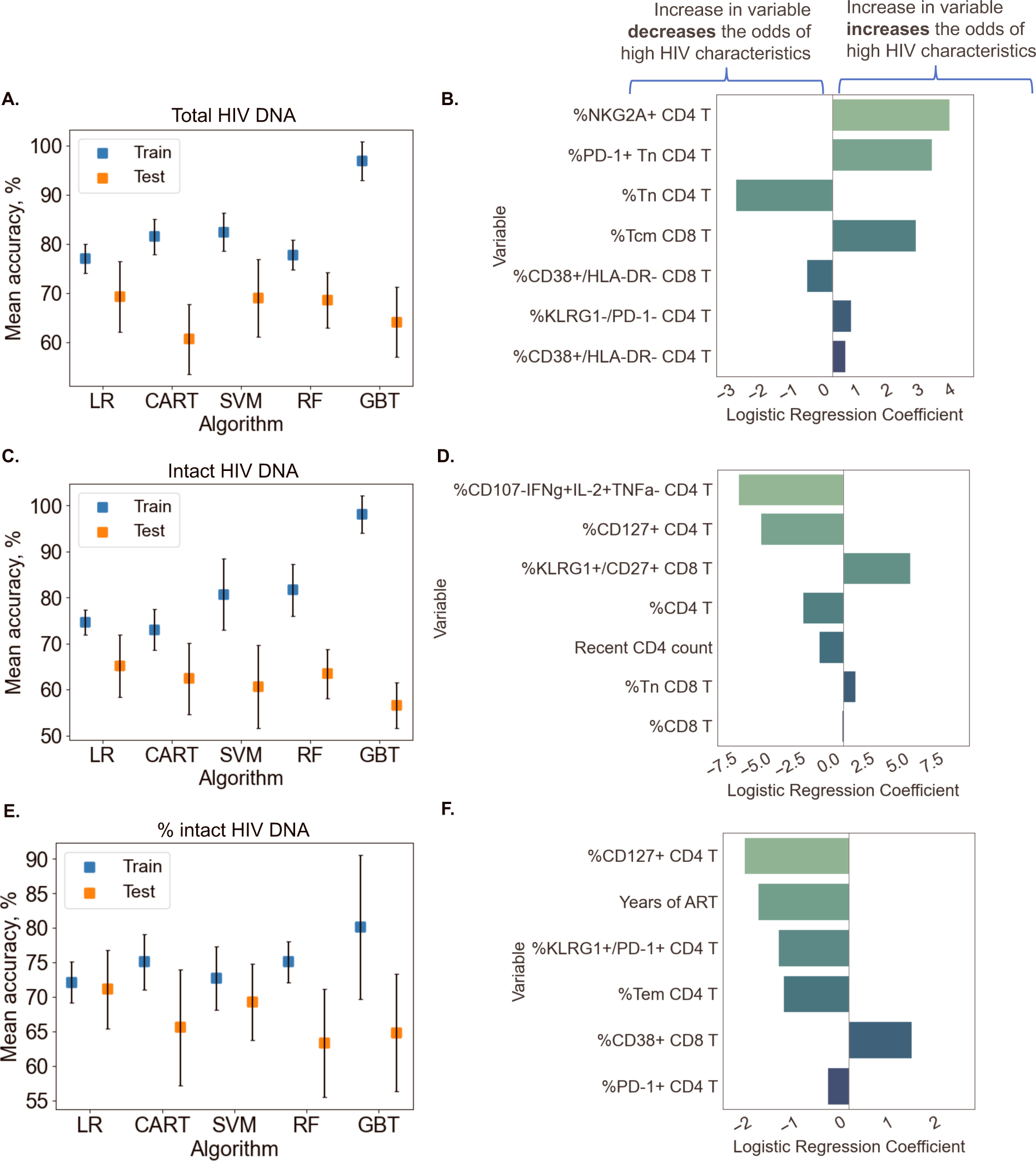
Predicting HIV reservoir characteristics with machine learning. Average training and test accuracies over ten training and test data splits for Random Forest (RF), Gradient Boosted Trees (GBT), Support Vector Machines with RBF kernel (SVM), Logistic Regression (LR), and CART models for total reservoir frequency (**A**), intact reservoir frequency (**C**), and percent intact (**E)** are shown. For one split of training and test sets, logistic regression models are visualized for total reservoir (**B**), intact reservoir (**D**), and percent intact (**F**). On the y-axis we show variables used by the model, while the x-axis displays coefficient values for individual variables used by models. For percent intact models we exclude participants with missing values of years of ART. For total and intact reservoir frequency, missing values of years of ART were imputed. The missing value of the CD4 nadir for one participant was imputed using the MICE algorithm.

To examine the contribution of individual immune features to model performance, we examined logistic regression coefficients for each immune cell variable in the model. Since Logistic Regression coefficients are associated with expected change in log odds (based on log*_e_*), we can think about the coefficient *β* for variable *X* in the following way: increasing variable *X* by one unit multiplies the odds of high reservoir frequency (probability that the reservoir size is high divided by the probability that the reservoir frequency is low) by *e^β^*. In **Figure 6B** we visualize the logistic regression model for one data split among ten we considered for total reservoir frequency. For this split we observe that higher values of %NKG2A+ CD4 T, %PD-1+ Tn CD4 T, and %Tcm CD8 T are associated with a increased probability of total reservoir frequency being high. On the other hand, an increase in %Tn CD4 T decreases the odds of high reservoir frequency. The model visualized in **Figure 6B** achieved 75.86% training and 75% test accuracy.

Similarly, logistic regression models performed best for predicting intact reservoir frequency compared to other methods (average 65.17% test accuracy, see **Figure 6C**). In **Figure 6D** we visualize the Logistic Regression model for one fixed data split. In this model we observed that higher values of %CD107a-IFN*γ*+IL-2+TNF*α*-CD4 T and %CD127+ CD4 T, and % CD4 T are associated with lower probability of intact reservoir frequency being high, while higher values of %KLRG1+CD27+ CD8 T are associated with increased probability of intact reservoir frequency being high.

For analysis of percentage intact HIV DNA, we display training and test accuracy values in **Figure 6E**. For the visualized model for one data split (**Figure 6F**), an increase in %CD127+ CD4 T, years of ART, %KLRG1+PD-1+ CD4 T, and %Tem CD4 T leads to a lower probability of percent intact HIV DNA being high.

Overall, these analyses demonstrate that we can use machine learning tools to construct models that can predict with approximately 70 percent accuracy whether a given PWH has qualitatively low or high total HIV DNA frequency, intact HIV DNA frequency or percentage of HIV DNA. Further studies with larger cohorts will likely improve the accuracy of these models.

## Discussion

In this cross-sectional study of peripheral blood samples from a well-characterized clinical cohort of 115 PWH on long-term ART, we used machine learning analysis of IPDA and high parameter flow cytometry to better understand the associations between the HIV reservoir and a broad range of immunophenotypes. Using these approaches, specific HIV reservoir features (total HIV DNA frequency, intact HIV DNA frequency, percentage intact HIV DNA) and clinical-demographic variables (age, sex, CD4 nadir, recent CD4 count, years on ART, years of HIV infection prior to initiating ART) were identified that predict specific immunophenotypes in PWH on long-term ART. Distinct host immune correlates of intact versus total HIV DNA frequency were also identified. Further, combinations of immune parameters that classified and predicted whether a given PWH has high or low total, intact, or percentage intact HIV DNA were defined. Collectively, this study confirms existing knowledge about HIV reservoir biology and identifies novel associations for further investigation. Additionally, from a methodological perspective, this work also demonstrates the utility of specific data science and machine learning approaches for HIV reservoir studies. Development and application of these analytic approaches are important to enable biological insights from high-parameter studies of the HIV reservoir and other complex biology, particularly with the advent of high-dimensional techniques such as CYTOF and scRNAseq.

Previous studies have sought to identify immune and other correlates of the total HIV DNA reservoir (21,26–44). These studies have identified several activation/exhaustion markers that correlate with total HIV DNA, including the CD4/CD8 ratio, T cell expression of HLA-DR, CD38, PD-1, CTLA-4, and LAG-3, and NK cell KLRG1 expression. However, not all studies have found associations of total HIV DNA and immune subsets, particularly for CD4 T cell HLA-DR and CD38 expression (39, 53–55) during longterm ART. There is also growing appreciation for additional factors that correlate with the HIV reservoir. Specifically, the size of the reservoir has been linked to the abundance of specific bacteria species in the intestinal microbiome (56). Also, genetic studies have indicated the existence of single nucleotide polymorphisms (SNPs) that are associated with reservoir size, including SNPs in MX1 and IRF7 (57–59). The correlative results from this present study corroborate many of these studies, and provide additional insights. A number of correlations between total HIV DNA and immune activation and exhaustion were identified. There were weak to moderate positive correlations observed between total HIV DNA frequency and CD4 T cell expression of NKG2A, PD-1, KLRG1 and HLA-DR as well as CD8 T cell expression of PD-1 and CCR7. Total HIV DNA frequency was inversely correlated with CD8 T cell expression of CD38, CD4 T cell expression of CD27, the frequency of naïve CD4 and CD8 T cells, and the CD4/CD8 ratio. Overall our results support the hypothesis that larger HIV reservoir size is associated with with increased frequencies of cells with an activated or exhausted phenotype, with depletion of naive cells and with an increased proportion of central memory CD8 T cells. The correlation of NKG2A expression in CD4 T cells with the total reservoir size represents a novel observation that merits further investigation. Only a very small percentage of CD4 T cells express NKG2A (less at 1 percent), and the biological significance for the correlation of these cells with the HIV reservoir is unknown. NGK2A is known to function as an inhibitory receptor for NK cells and for CD8 T cells, but its role in CD4 T cells is less clear (60).

When we examined immune correlates of intact HIV DNA measured by the IPDA, many of the same relationships as described above were noted, but with decreased magnitude and often lacking statistical significance. This is consistent with a recent study using the IPDA that did not find associations of intact HIV DNA with CD4 T cell HLA-DR and CD38 co-expression (61), as well as a study using near-full-length viral sequencing in single cells that found only slight enrichment of HLA-DR or PD-1 in cells with genetically intact, inducible proviruses (36). The stronger relationship between the immune cell phenotypes and the total reservoir, rather than the intact reservoir, should be investigated further. We speculate that this association could be driven by the more numerous defective proviruses promoting inflammation or immune activation through innate immune sensing pathways, or a greater level of transcriptional activity for defective proviruses. This is certainly plausible giving the growing appreciation for the immunogenicity and high frequency of defective proviruses within the HIV reservoir (62–65). It is important to note, however, that it is also possible that the stronger association of total proviral burden with immunophenotype reflects persistent damage caused to the immune system before therapy, and that this association between the total reservoir and immune cells may not result from ongoing interactions between HIV and the immune system during therapy.

Uniquely, CD127 expression on CD4 T cells was significantly associated with the intact reservoir frequency but not the total reservoir. Specifically, the frequency of CD127-CD4 T cells was positively correlated with the frequency of intact proviruses. CD127 is the alpha-chain IL-7 receptor and is expressed on long-lived memory T cells that are depleted during untreated HIV infection. Post-ART, the frequency of CD127+ memory CD4 subsets in PWH increases over time, albeit slowly (11). Interestingly CD127+ cells are enriched in latent HIV infection in tissues (66), and IL-7 is associated with slower natural reservoir decay (27). Furthermore, IL-7 signaling is known to drive expression of VLA-4, which was recently reported to highly enrich for genetically intact, inducible proviruses (36). This observational finding that intact but not total HIV DNA inversely correlates with CD127 expression on CD4 T cells requires further investigation.

Dimension reduction machine learning approaches identified two robust clusters of PWH when using total HIV DNA reservoir-associated immune cell frequencies. These clusters exhibited distinct reservoir characteristics, with one cluster being enriched with participants with a larger total reservoir size, a lower percentage of intact viruses, a lower frequency of naïve T cells and elevated expression of activation and exhaustion markers, while the other contained PWH with smaller total reservoirs, higher percentage of intact viruses, higher frequency of naïve T cells and lower expression of activation and exhaustion markers. The cluster membership was notably associated with age and time on therapy. The existence of two distinct clusters of PWH with different immune features and reservoir characteristics could have implications for HIV cure strategies. These two clusters are also interesting in the context of a previous study examining reservoir characteristics which proposed two major types of viral reservoir within PWH: one type with smaller reservoirs that were enriched in Tcm cells and associated with lower levels of Ki67+ cells and immune activation, and another type with larger reservoirs that were enriched in Ttm cells and associated with higher levels of Ki67+ cells and immune activation (27). It is possible that these two proposed types of reservoirs represent the two clusters we observe using immune markers and dimension reduction, although we were unable to examine the frequency of HIV proviruses specifically within Tcm and Ttm cells in this study.

In addition to unsupervised dimension reduction approaches, we were able to construct simple, in-terpretable decision trees that describe our cohort of PWH with more than 80% accuracy with respect to the size of the reservoir. Furthermore, when we constructed decision trees to classify PWH separately based on total reservoir frequency, intact reservoir frequency and percentage intact, distinct sets of features were important (though some were the same) for each of these models. The decision tree analyses highlight complex immune parameter combinations that serve as a basis for unique hypotheses about HIV reservoir biology. One striking example of machine learning approaches identifying potential reservoir biology is the percentage intact HIV DNA decision tree. As expected, years of ART was critical to predict the percentage of intact proviruses, but CD4 T cell CD127 expression, CD4 T cell KLRG1 expression and recent CD4 count also played a significant role.

Logistic regression models that we trained to classify high versus low total reservoir, intact reservoir, and percentage intact were able to predict out-of-sample with *∼*70% accuracy on average. Each of these models relied on five features only and were simple enough for us to visualize and assess how important each variable is for every characteristic of reservoir size. These models corroborated the potential biological relationships identified in the decision tree analysis, and confirmed that these variables could actually be used to predict out-of-sample whether total, intact or percentage intact HIV DNA was qualitatively high or low. The success of our predictive models was likely limited by the relative small size of the datasets used to train and test them (n=115). As such, larger datasets will be required to generate more accurate models that can be used for reliable prediction of reservoir characteristics. Additionally, it is likely that unmeasured factors and biological noise also contribute to reservoir size.

Our findings should be considered in the light of some inherent limitations and caveats. This study is cross-sectional in nature and is primarily observational, so the findings should be interpreted with caution. As with any observational study, confounding bias may influence our results, particularly for correlative analyses. The associations between the HIV reservoir and the immune system we observed were fairly weak, despite being statistically significant. Similarly, the predictive ability of our ML models was modest (70 percent) for classifying PWH as having an above or below median reservoir. As mentioned above, our cohort is likely too small to generate accurate predictive ML models, and consists of mostly male participants, which could bias the results. We also cannot rule out the influence of additional unmeasured confounding factors. In particular, the length of untreated infection prior to ART could be a major contributor to the immune signatures we observe. In this study, the participants had all been treating during chronic infection, and different associations might be observed in PWH who are treated during acute infection. Another limitation to this study is that our analyses only considered cells and proviruses in the peripheral blood, and this may not represent HIV reservoirs and immune cells located in tissues known to harbor latent HIV such as gut, spleen and brain. Our analysis of surface protein phenotypes also does not detect potentially important associations between the reservoir and transcriptional pathways within these cell types. Future work using single cell RNAseq from a similar cohort may help to reveal deeper layers of associations that are invisible to the current analysis. As is always the case with correlation studies, its is important to be aware that these associations are only correlations and do not necessarily represent functional or mechanistic relationships. Even so, these correlations are still useful as a source of novel hypotheses that could be tested in subsequent studies.

Nevertheless, our study does have some strengths compared to previous work in this area, specifically a highly detailed flow panel, allowing us to quantify numerous immune subsets with a high degree of resolution. Also, the use of the intact proviral DNA assay allows us to separately examine the association of the immune system with intact and total viral DNA. It is however, important to note that IPDA does not measure all proviral sequences in PWH and is likely failing to identify some sequences due to proviral polymorphisms or dual deletions in the two IPDA amplicons. Also, a minor fraction of IPDA+ proviruses are actually defective due to the limited specificity of using two short amplicons to select for full-length intact proviruses (46, 67).

Overall, these findings suggest a complex concert of immune recovery, HIV reservoir dynamics, and intrinsic host factors (age, biological sex) that shapes host immunophenotype, even after years of ART. Mechanistic work will be needed to fully dissect the dynamic relationship between the immune system and the reservoir. Nevertheless, these findings support a model in which ongoing interaction between the HIV reservoir and the host immune cells continue to drive an association, albeit a relatively minor one, with persistent CD4 T cell activation during long-term ART. Notably, both intact and defective proviruses appear to contribute to this immune signature of HIV latency during long-term ART. This study identifies several areas for future investigation. The question of whether the HIV reservoir directly drives persistent immune activation could be addressed directly by specifically suppressing viral transcription during therapy and examining the impact on the immune system. The stronger association of the intact reservoir with CD127 expression in CD4 T cells is an intriguing observation that warrants further validation and explanation. This observation could, for example, be explained by a model in which the intact reservoir is preferentially located in cells with a distinct pattern of CD127 expression. These ideas, along with other future studies exploring the interplay of the host immune system and HIV reservoir might leverage new high-dimensional technologies such as scRNAseq and mass cytometry. The machine learning approaches described and applied in this study may be particularly useful for gaining biological insights into these high parameter datasets.

## Methods

### Cohort and sample collection

115 people with HIV (PWH) were recruited from two clinical sites. 66 PWH were recruited at Duke University medical center and 49 PWH at UNC Chapel Hill. Peripheral blood was collected and peripheral blood mononuclear cells (PBMCs) were obtained by Ficoll separation, then frozen in 90% fetal bovine serum 10% dimethylsulfoxide (DMSO) (51, 68–70).

### Flow cytometry

Cell preparation and staining followed previously described methods (71). All antibodies were titrated to optimize signal-to-noise ratio on PBMCs prior to use, assuming a 50μL staining volume. Cell viability was examined using Zombie-NIR Fixable Viability Dye (0.4μL per 50μL staining volume; Biolegend). Antibodies used for surface staining were as follows: KLRG1-BV421 (SA231A2 clone, Biolegend, CD45RA-PacBlue (H100 clone, BL), CD8-BV570 (RPA-T8 clone, Biolegend), CD127-BV605 (A019D5 clone, Biolegend), CD56-BV650 (5.1H11 clone, Biolegend), CCR7-BV711 (G043H7 clone, Biolegend), CD27-BV750 (O323 clone, Biolegend), PD1-VioBright515 (REA165 clone, Miltenyi), NKG2A-PE-Vio615 (REA110 clone, Miltenyi), CD16-PerCP-Cy5.5 (33G8 clone, Biolegend), CD38-PCPeF710HB7 clone, TF), CD14-SparkNIR685 (63D3 clone, Biolegend), CD19-SparkNIR685 (HIB19 clone, Biolegend), HLA-DR-APC-F750 (L243 clone, Biolegend). Antibodies used for intracellular staining were as follows: CD3-BV480 (UCHT1 clone, BD), CD4-PerCP (L200 clone, BD), IFN-g-PE-Cy7 (4S.B3 clone, Biolegend), IL-2-APC (MQ1-17H12 clone, Biolegend), TNFα-AF700 (Mab11 clone, Biolegend). For HIV peptide stimulation, the cells were incubated with peptide mixes from Gag, Pol and Env proteins at 0.2*µ*g/mL in the presence of brefeldin A and monesin for 6 hrs. Samples were analyzed using a Cytek Aurora spectral flow cytometer. Flow cytometry analysis was performed in FlowJo v10.8 software.

### Intact Proviral DNA Assay (IPDA)

Cryopreserved samples of PBMCs from each study participant were viably thawed. A portion was used for immunophenotyping as described above, and the remainder were subjected to total CD4 T cell negative selection with the StemCell Technologies EasySep^TM^ Human CD4+ T Cell Enrichment Kit (Cat# 19052). CD4 T cell DNA was extracted using the QIAamp DNA Mini Kit and quantified on a NanoDrop 1000 (Thermo Fisher Scientific). IPDA was performed as originally described (46), with a validated PCR annealing temperature modification to increase signal to noise ratio (70). Gating for positive droplets was set using negative (DNA elution buffer and HIV-seronegative CD4 T-cell DNA), and positive (Integrated DNA Technologies gblock amplicon) positive control wells processed in parallel. DNA shearing index values were similar to those reported previously (median, 0.33; interquartile range [IQR], 0.31– 0.34) (68–70). A median of 1.04*x*10^6^(*Q*1 − 8.71*x*10^5^*, Q*3 − 1.16*x*10^6^) cell equivalents were assessed for each donor.

## Statistics

### Variable data analysis

To create binarized labels that represent reservoir characteristics, we split the data at the median, which is 553/M for total reservoir frequency, 53/M for intact reservoir frequency, and 8.64 percent for percent intact. For every variable (immune cell frequency, demographics, and clinical information), we computed Spearman correlation and an Area Under the Curve (AUC) value. To compute the AUC value, we first created a Receiver Operating Characteristic (ROC) curve by plotting the true positive rate versus the true negative rate for every cell subset frequency. AUC is then computed as the area under the ROC curve using the trapezoidal rule.

To assess the importance of clinical, demographic information and HIV reservoir characteristics in describing the immune markers, we performed a leave-one-covariate-out (LOCO) analysis. We trained a linear regression model (*M*_1_) with intercept based on all variables. We removed each variable one at a time and trained a new model without this variable. Then we computed the difference in R^2^ score before and after dropping the variable. Variance Inflation Factors (VIFs) for age, sex, years of ART, CD4 nadir, recent CD4 account, and years of HIV infection prior to HIV infection were less than 2, indicating an acceptable level of correlation between these independent variables (72). However, higher VIF values were observed for intact and total HIV DNA, therefore separate LOCO inference analyses were conducted in order to avoid artificial fluctuations in model fit due to multicollinearity.

### Data visualization

We used PaCMAP (50) to reduce the dimensionality of the dataset to a 2-dimensional space. Due to the relatively small sample size of our dataset (n=115), we set the number of neighbors to 5. Additionally, we used GOSDT (52) which computes sparse optimal trees to identify patterns in the dataset. For all the visualization trees we set the depth budget parameter to 4 and the regularization parameter to 0.02.

### Training and generalization

Our training procedure for classification is described in **Algorithm 1** in Supplemental Materials, which is a standard machine learning pipeline with an added search for the number of variables. For the full hyperparameters list see **Table S8**. We normalized continuous variables to have values between 0 and 1. For logistics regression models, we set the regularization parameter of *ℓ −* 2 regularization based on cross-validation. The values are 10 for total reservoir frequency and percent intact and 100 for intact reservoir frequency.

## Study approval

Written informed consent was obtained for all study participants. The study design was reviewed and approved by IRB for both Duke University and UNC Chapel Hill.

## Data availability

Underlying data available at: https://doi.org/10.15139/S3/3CFVIU.

Code is available at: https://github.com/lesiasemenova/ML_HIV_reservoir.

## Author contributions

Designed the study – EPB, DMM, CDR. Conducted experiments – SDF, ADVC. Analyzed the data – LS, YW, SDF. Wrote the manuscript: All authors.

## Acknowledgements.

This work was supported by the following grants from the National Institutes of Health: NIAID R01 AI143381 (EPB), NIAID UM1 AI164567 (DM Margolis), NIDA R61 DA047023 (EPB), NIDA R01 DA054994 (CDR), NIAID F30 AI145588 (SDF).

## Supplementary Methods

### CD4/CD8 ratio regression analysis

To examine the connection between CD4/CD8 and (CD127+ CD4)/CD8 ratio and HIV characteristics (**Figure S5**), we first computed the natural logarithm (log_e_) of the intact, total and percent intact as the outcome and normalized outcome of each immune feature (meaning that our values of ratio and outcome are between 0 and 1). We removed outliers using the DBSCAN algorithm (1), which is an unsupervised clustering algorithm that groups data points based on density into a single cluster. For DBSCAN, we set the parameter that determines radius of a circle around each point (that is used to compute density) as 0.15 and minimum number of samples in each cluster as 10. Then we fitted the remaining datapoints using Linear Regression.

### Regression machine learning models to predict reservoir frequency from immune cell frequencies

To predict reservoir frequency based on immune cell frequencies, clinical and demographic information (**Figure S13**), similar to classification, we sought to find the set of model parameters and types of models that would be able to fit the dataset without overfitting. We used several approaches: Linear Regression (LR), Ridge Regression (RR), Kernel Regression with RBF kernel (KR), Decision Tree Regressor (DT), Random Forest (RF), and Gradient Boosted Trees (GBT). We followed the procedure described in Algorithm 1, optimized the R^2^ score, returned the mean and standard deviation of the R^2^ score of training and test sets, and chose features based on the absolute value of the correlation coefficient. We normalized variables to have values between 0 and 1. We then fitted the models to the normalized natural logarithm (log_e_) of total, intact reservoir frequency and percent intact and then further normalization makes outcomes to be between 0 and 1. For total and intact reservoir frequency we visualized the Ridge Regression model, which performed the best based on mean test *R*^2^ score (**Figure S13 D, E, G, H**). We set the regularization parameter to 1, as this value was best on average according to cross-validation for total and intact reservoir frequency. For percent intact, we visualized Linear Regression based on 4 variables (**Figure S13 F, I**) as it achieved the second highest *R*^2^ score after Kernel Ridge based on 25 variables.

---

**Figure S1:**
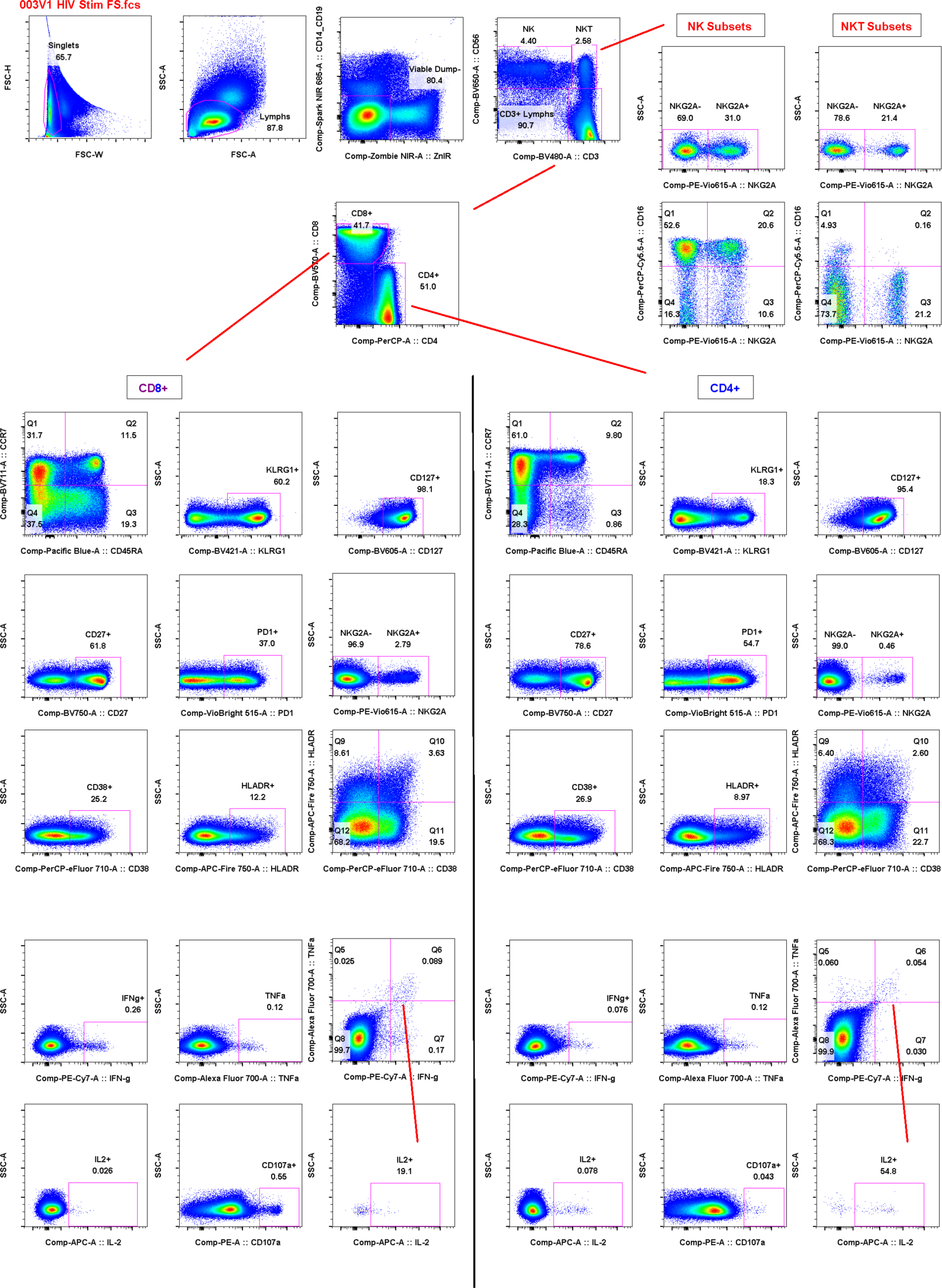
Representative flow cytometry gating is shown for one sample from the 115-person cohort.

**Table S1:** Intact, Total reservoir frequency and % intact for demographic subgroups. We report medians and [Q1, Q3] for total reservoir frequency, intact reservoir frequency (per million CD4 T cells), and percent intact. Mann-Whitney U test was used to compute p-values for gender and Kruskal-Wallis H-test for race.

| **Group** | **Size** | **Intact HIV DNA** | **Total HIV DNA** | **% intact HIV DNA** |
| --- | --- | --- | --- | --- |
| **Male** | 88 | 63.5 [18.75 146.25] | 681 [198 1433.75] | 8.73 [4.9 20.58] |
| **Female** | 27 | 41 [19, 92.5] | 489 [317.5, 637] | 8.64 [4.86, 15.56] |
| **p-value** |  | 0.4521 | 0.1937 | 0.9451 |
| **Black** | 69 | 43 [19, 113] | 502 [181, 1246] | 9.77 [4.64, 16.4] |
| **White** | 43 | 69 [26 191.5] | 605 [289.5 1415.5] | 8 [5.31 20.81] |
| **Other** | 3 | 67 [41, 520] | 1055 [720, 1165,] | 6.39 [5.20, 41.37] |
| **p-value** |  | 0.5079 | 0.4916 | 0.9914 |

**Table S2:**
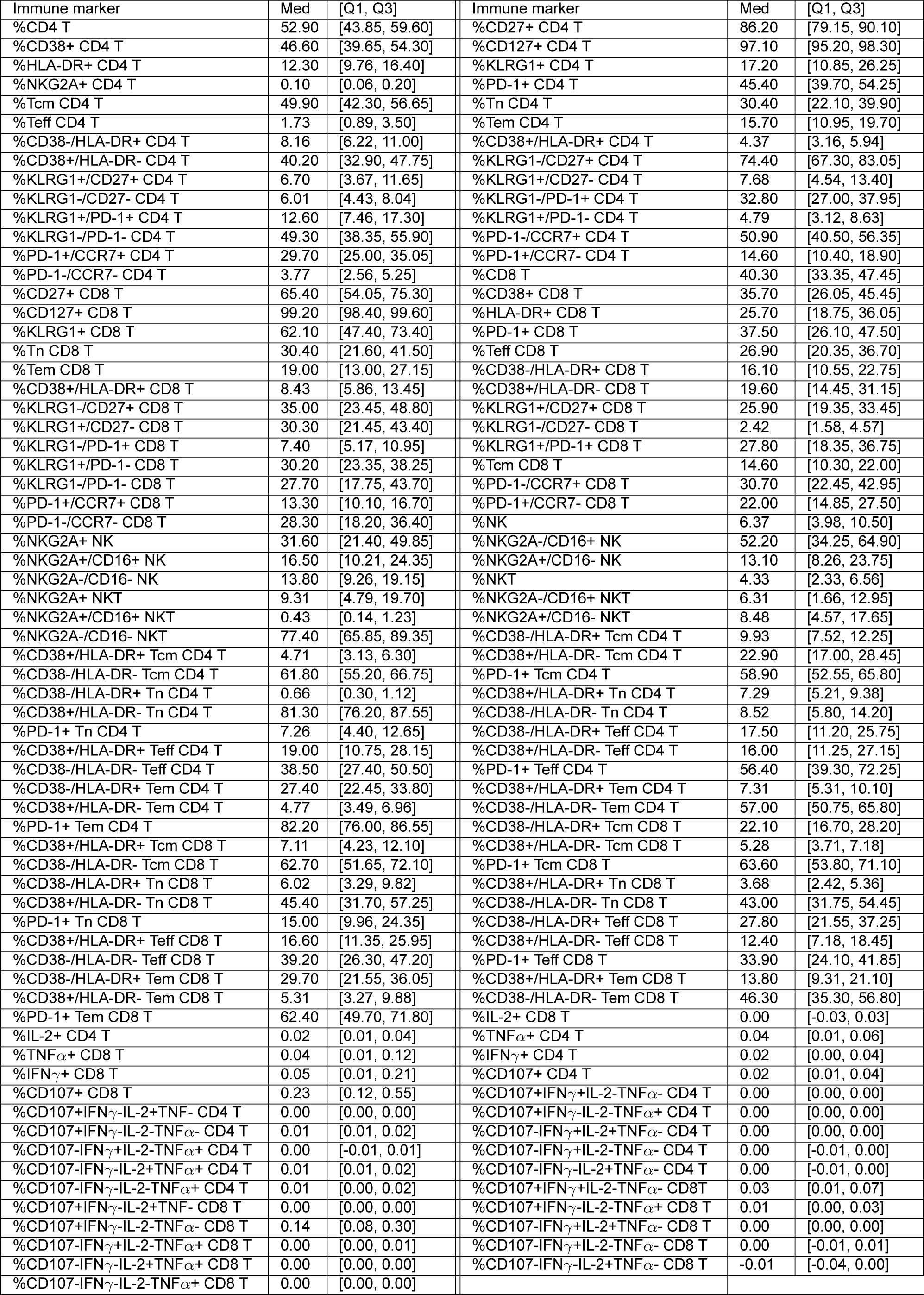
Immune subsets characteristics. . We report medians, 25% and 75% percentiles for every immune variable used in the study.

**Table S3:**
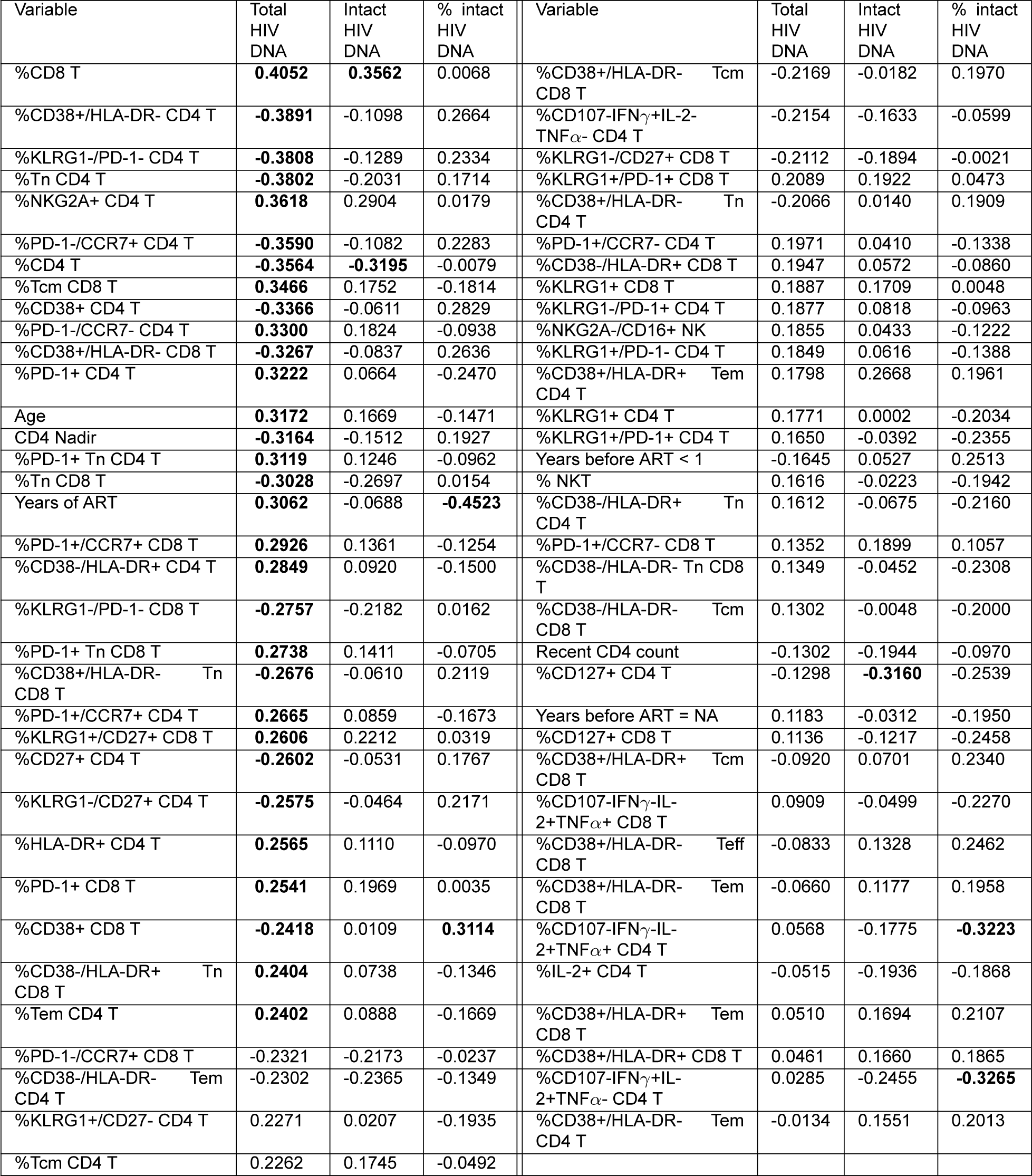
Host features correlate with HIV reservoir characteristics. The abundance of 133 immune cell populations was determined by flow cytometry and the HIV reservoir was quantified by intact proviral DNA assay for a cohort of 115 people with HIV (PWH). Each abundance and clinical and demographic variable was correlated with total HIV reservoir frequency, intact reservoir frequency, and the percentage of intact proviruses. Spearman correlation coefficients are shown for 69 variables that were correlated (p*<*0.05) with one or more reservoir characteristics. Each feature/subset is ranked by the absolute value of the correlation coefficient for the total reservoir frequency. Variables that had significant p-values (*<*0.05) after Benjamini–Hochberg correction for multiple comparisons (144 comparisons in our case) with one of the characteristics are shown in bold. For years of ART, correlations are computed based on 108 participants, excluding participants with missing years of ART values. For CD4 Nadir, correlations are computed based on 114 patients, excluding participants with the missing CD4 Nadir value.

**Figure S2:**
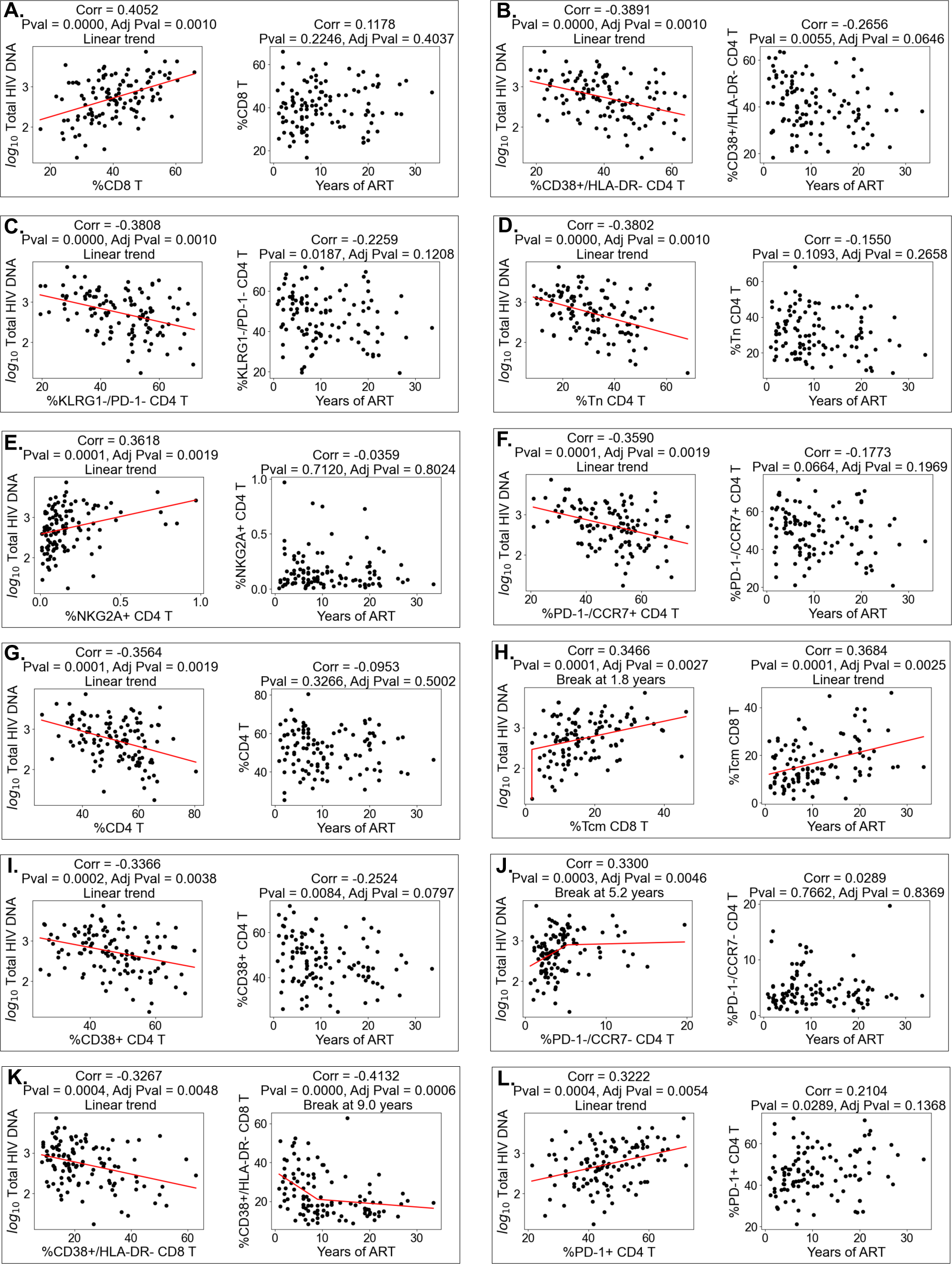
Abundance of immune cell subsets correlates with HIV reservoir. Continuation in Figures S3, S4.

**Figure S3:**
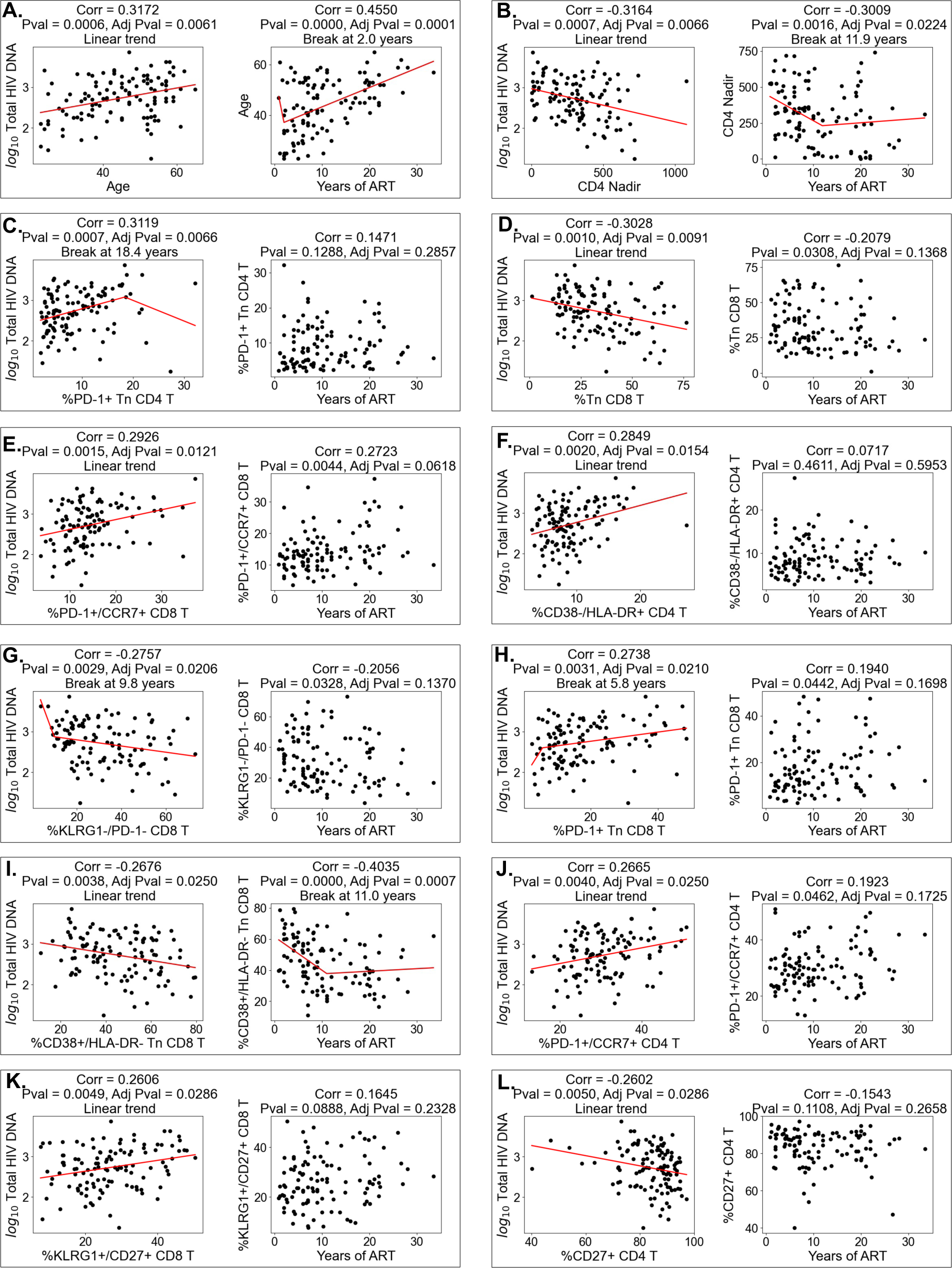
Abundance of immune cell subsets correlates with HIV reservoir. Continuation in Figures S2, S4.

**Figure S4:**
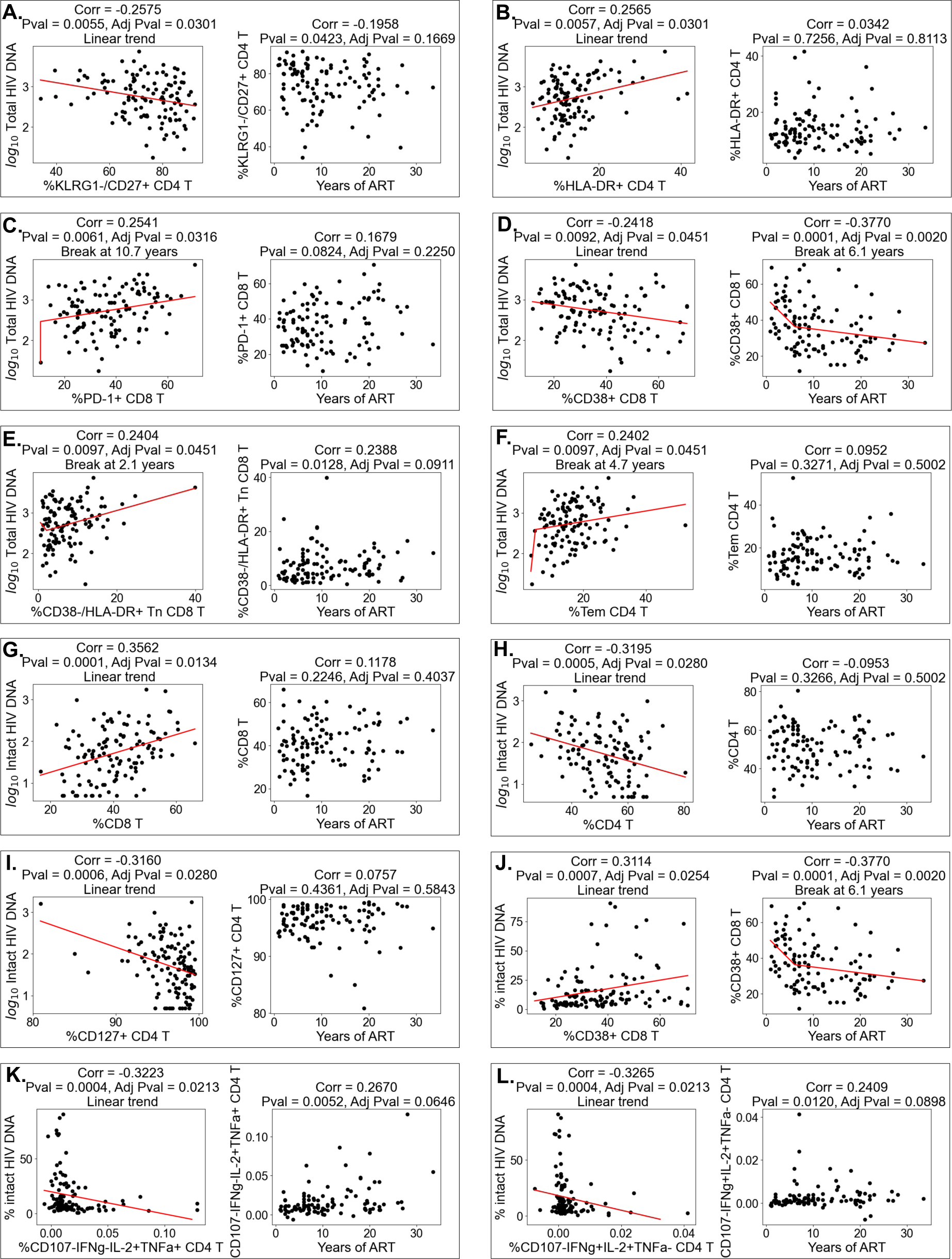
Abundance of immune cell subsets correlates with HIV reservoir. Continuation in Figures S2, S3.

**Figure S5:**
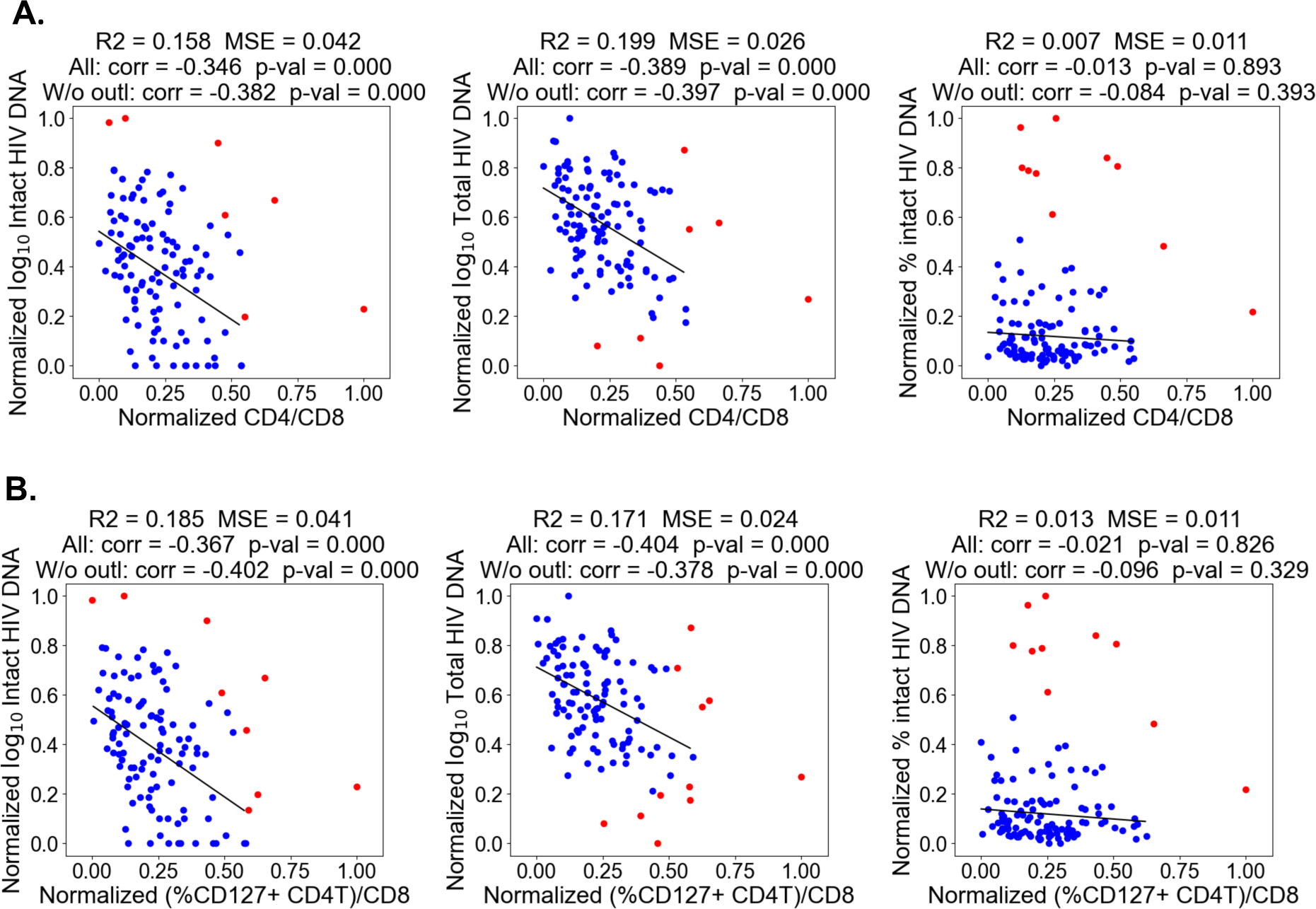
CD4/CD8 and (%CD127+ CD4T)/CD8 ratios correlate with total and intact reservoir frequency. Normalized (values transformed to be between 0 and 1) CD4/CD8 (**A**) and (%CD127+ CD4T)/CD8 (**B**) ratios are shown on the x-axis and the normalized natural logarithm (log_e_) of the total reservoir, intact reservoir and percent intact on the y-axis. Spearman correlation was computed between ratios and HIV reservoir characteristics. Outliers (red data points) were removed with the DBSCAN clustering algorithm and a linear regression model was fitted (black line) to the remaining data points. Spearman correlations, R^2^ scores and mean squared error after outlier removal are displayed.

**Table S4:** PWH immune features correlate with years of ART. The abundance of 133 immune cell populations was determined by flow cytometry and the HIV reservoir was quantified by intact proviral DNA assay for a cohort of 115 people with HIV (PWH). Each abundance and clinical and demographic variable was correlated with years of therapy for all participants (middle column) and for participants who had been on therapy for less than 10 years (right column). Spearman correlation coefficients are shown for all variables that were correlated (p*<*0.05) after Benjamini-Hochberg correction with years of treatment for all participants or for participants that were on ART for less than 10 years. Variables that had significant p-value (*<*0.05) after correction for multiple comparisons are shown in bold. Correlation is computed based on 108 PWH for all participants and on 60 PWH for PWH who were on therapy for less than 10 years). Computations exclude PWH with missing years of ART values. For CD4 Nadir, correlations are computed excluding the participant with missing CD4 Nadir value.

| Variable | Spearman correlation coefficient,<br>All participants | Spearman correlation coefficient,<br>Participants with years of ART < 10 years |
| --- | --- | --- |
| Age | **0.4550** | 0.2629 |
| %CD38+/HLA-DR- CD8 T | **-0.4132** | -0.3788 |
| %CD38+/HLA-DR- Tn CD8 T | **-0.4035** | -0.3075 |
| %CD38+ CD8 T | **-0.3770** | -0.3025 |
| %Tcm CD8 T | **0.3684** | 0.1595 |
| %CD38-/HLA-DR- Tn CD8 T | **0.3580** | 0.2222 |
| %CD38+/HLA-DR- Tcm CD8 T | **-0.3138** | -0.2666 |
| CD4 Nadir | **-0.3078** | 0.1265 |
| %CD107a+IFNγ-IL-2+TNF- CD8 T | **0.2903** | 0.1728 |

**Table S5:** Multicolinearity analysis for variables used in models to predict immunophenotypes. . Variance Inflation Factor (VIF) for every variable is computed for models that fit immunophenotypes as a target and are based on clinical, demographic information and HIV characteristics (total, intact reservoir frequency or percent intact) as features. Note that here “Years before ART” is a continuous variable that we used for VIF computations only. In our other analysis we categorize this variable and use “Years before ART=NA”, “Years before ART*<* 1”, and “Years before ART*≥* 1” instead. We excluded 3 participants with race indicated as “Other” to avoid near-binary categorical variables in VIF computation.

| Total HIV DNA model |  | Intact HIV DNA model |  | % intact HIV DNA model |  |
| --- | --- | --- | --- | --- | --- |
| Variable | VIF value | Variable | VIF value | Variable | VIF value |
| Total HIV DNA | 1.2545 | Intact HIV DNA | 1.0517 | % intact HIV DNA | 1.3469 |
| Age | 1.7426 | Age | 1.7243 | Age | 1.7220 |
| Sex | 1.1948 | Sex | 1.1616 | Sex | 1.1795 |
| Race | 1.5080 | Race | 1.4577 | Race | 1.6348 |
| Years of ART | 1.3048 | Years of ART | 1.3004 | Years of ART | 1.3181 |
| CD4 nadir | 1.3188 | CD4 nadir | 1.2875 | CD4 nadir | 1.3773 |
| Recent CD4 count | 1.3314 | Recent CD4 count | 1.3374 | Recent CD4 count | 1.3134 |
| Years before ART | 1.1367 | Years before ART | 1.1138 | Years before ART | 1.1611 |

**Figure S6:**
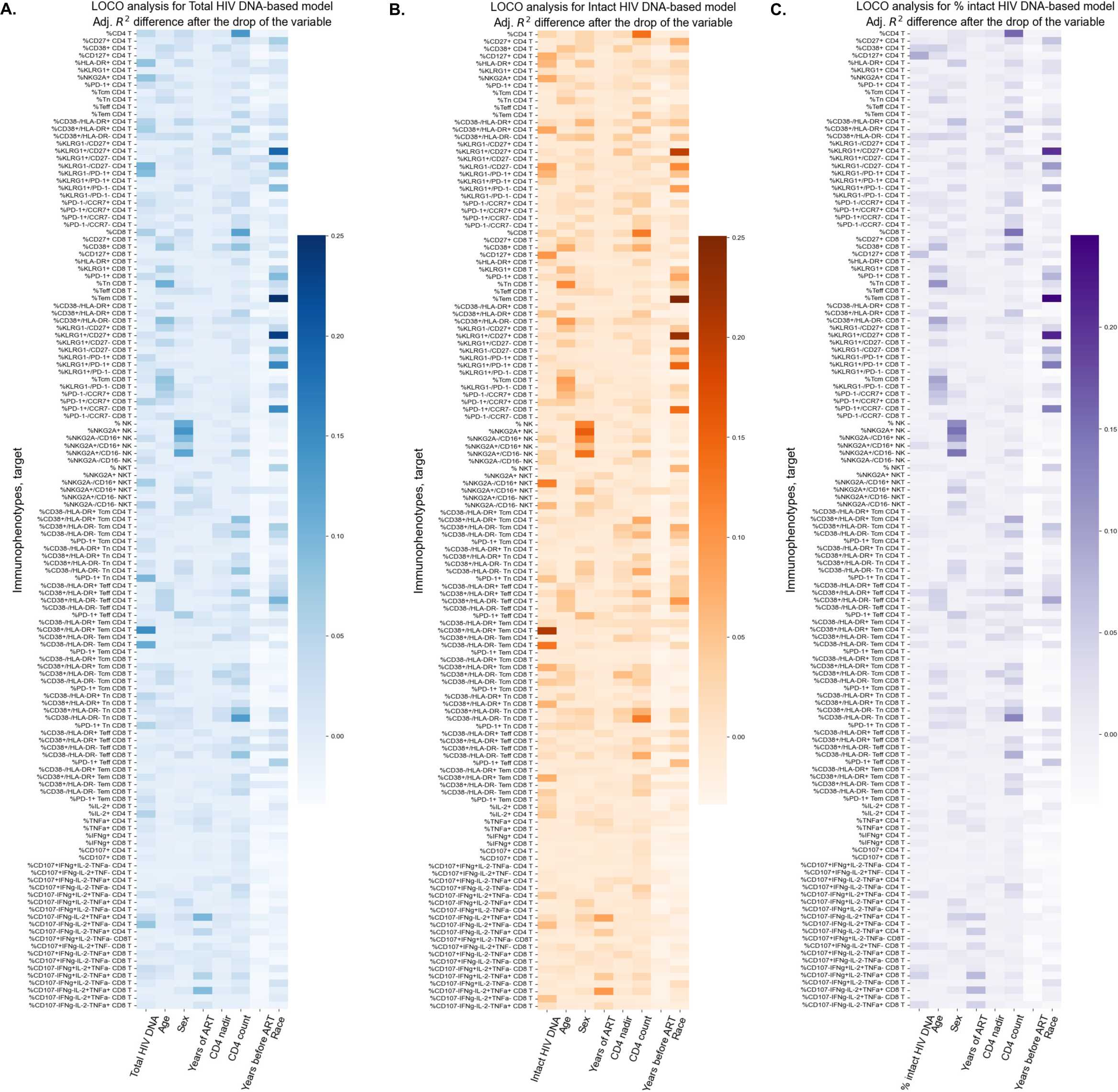
Leave-one-covariate-out (LOCO) analysis for clinical-demographic features and reservoir characteristics while predicting immunophenotypes. Analysis that is described in **Figure 2** was performed for all 133 immunophenotypes considered in the study. Drops in adjusted R^2^ scores are reported for models that use total reservoir frequency (**A.**), intact reservoir frequency (**B.**), or percent intact (**C.**) as features in addition to clinical and demographic information such as age, biological sex, race, years of treatment, CD4 nadir, recent CD4 count, and years of HIV before treatment (=NA, *<* 1, *≥* 1). On the x-axis, we show features that were dropped from the model. On the y-axis, we display immunophenotypes, which are targets (outcomes) for the linear regression models. In **Tables S6-S8** we show the actual values of drops in adjusted R^2^ score.

**Figure S7:**
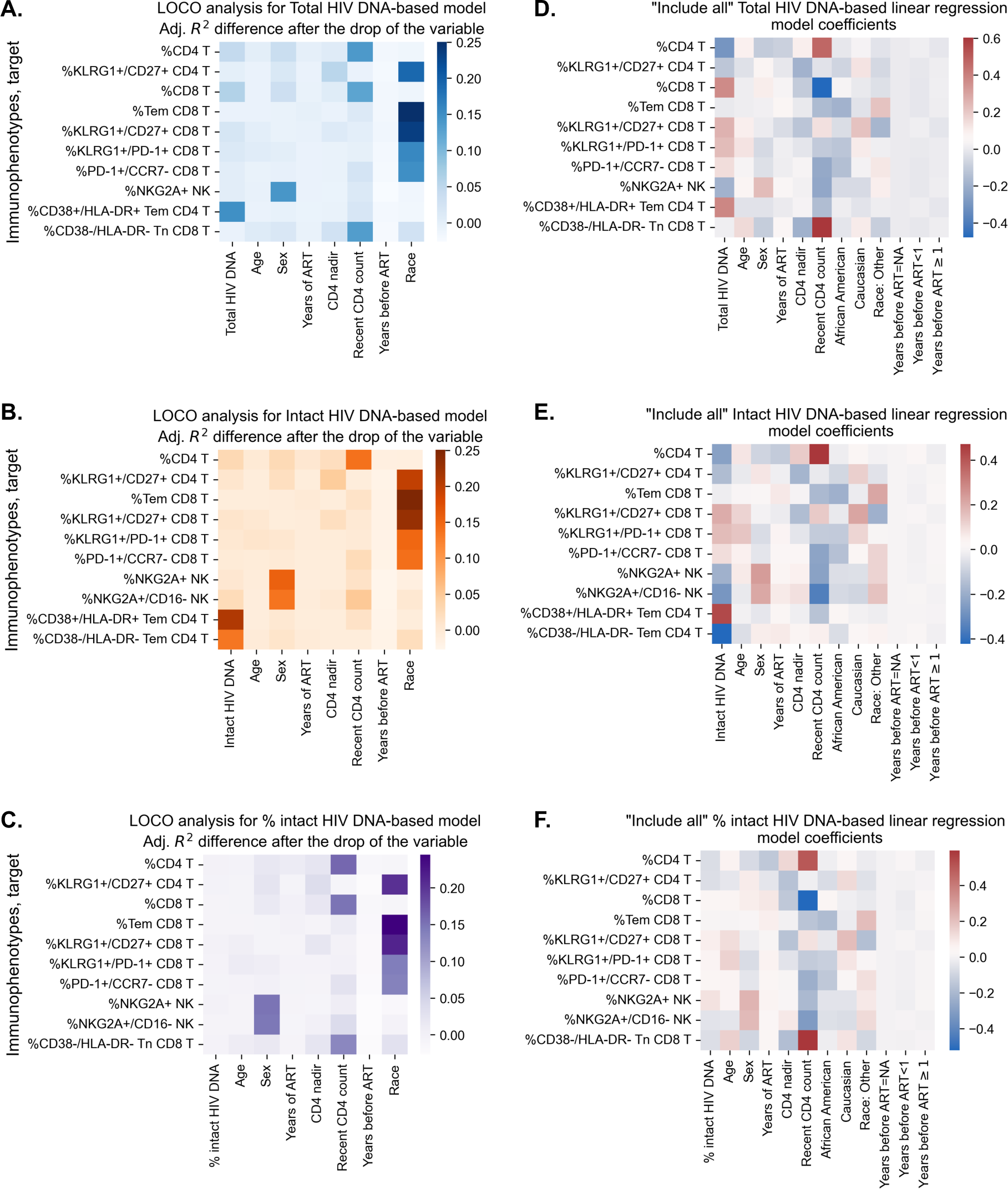
Coefficient visualization for linear regression models that predict immunophenotypes based on clinical, demographic information and HIV characteristics. **(A., B., C.)** LOCO analysis from **Figure 3** for total-reservoir based (**A.**), intact reservoir-based (**B.**) and percent intact-based (**C.**) models. The drops in adjusted R^2^ scores are shown after removing a feature and training a new model without it. **(D., E., F.)** Coefficient visualization for models that include clinical and demographic information such as age, biological sex, years of treatment, CD4 nadir, recent CD4 count, and years of HIV before treatment = NA, years of HIV before treatment *<* 1, years of HIV before treatment *≥* 1 and total reservoirs frequency (**D.**) or intact reservoir frequency (**E.**), or percent intact (**F.**). No features are dropped from these models, they are “Include all” models from tables **Tables S6-S8**. On the x-axis, the feature is shown, and on the y-axis the target (immunophenotypes from (A., B., C.)). The heatmap displays the coefficient in front of that variable in the model (if the model is %CD4 T= *β*_1_Total + *β*_2_Age + *β*_3_Sex + *…*, then *β*_1_*, β*_2_*, β*_3_*, …* are visualized), where positive coefficients are shown in red and negative in blue.

**Table S6:**
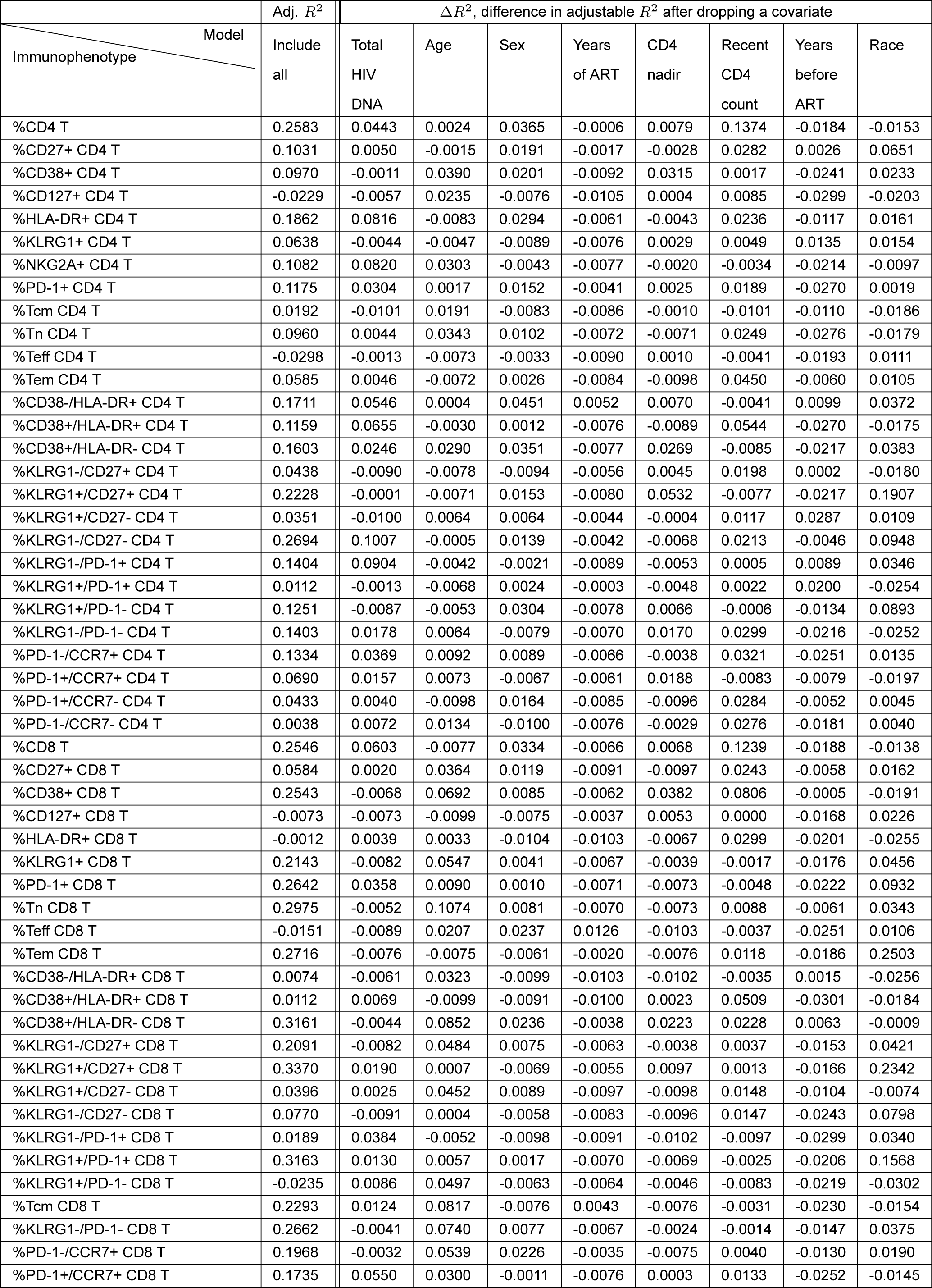
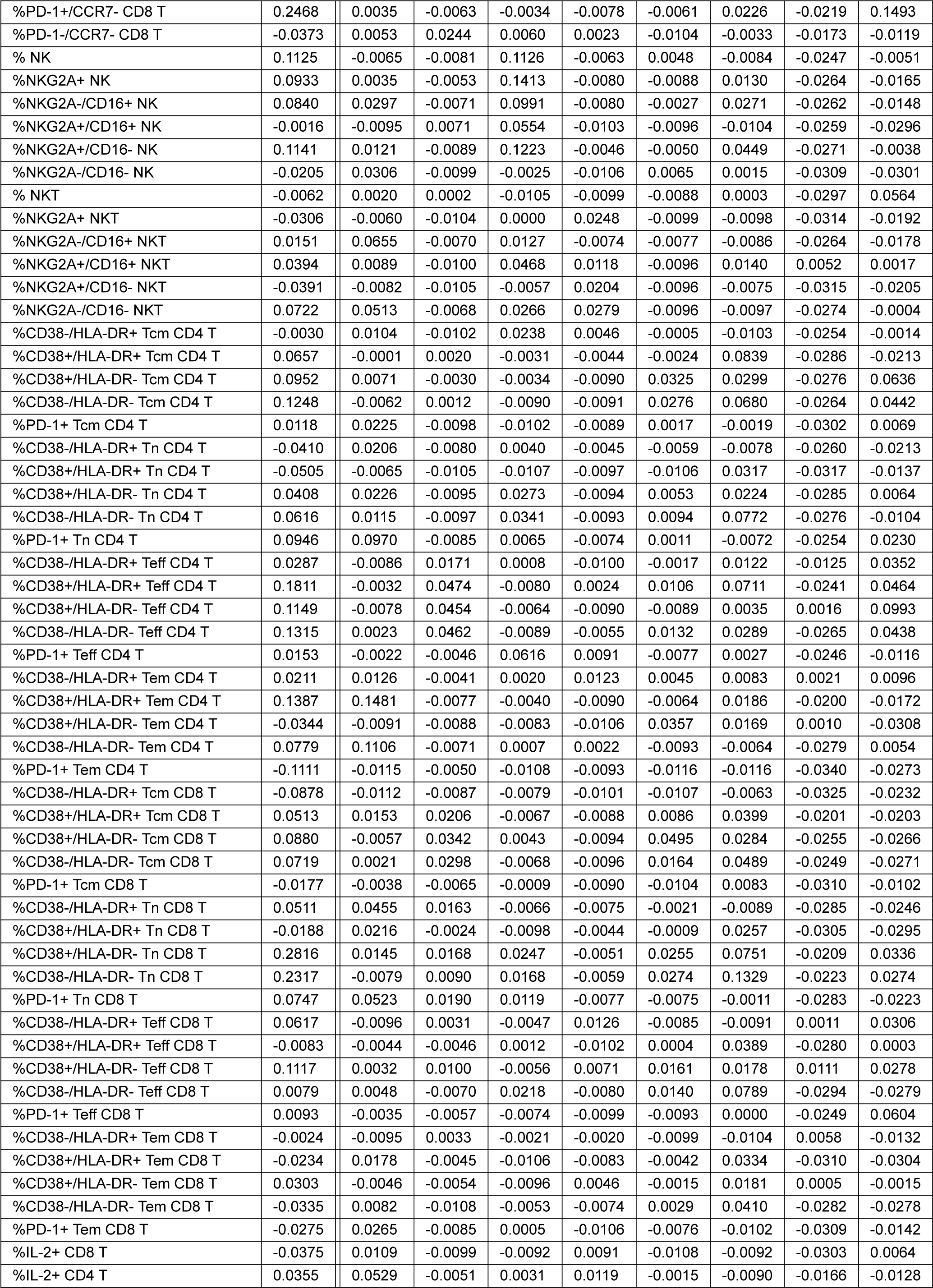
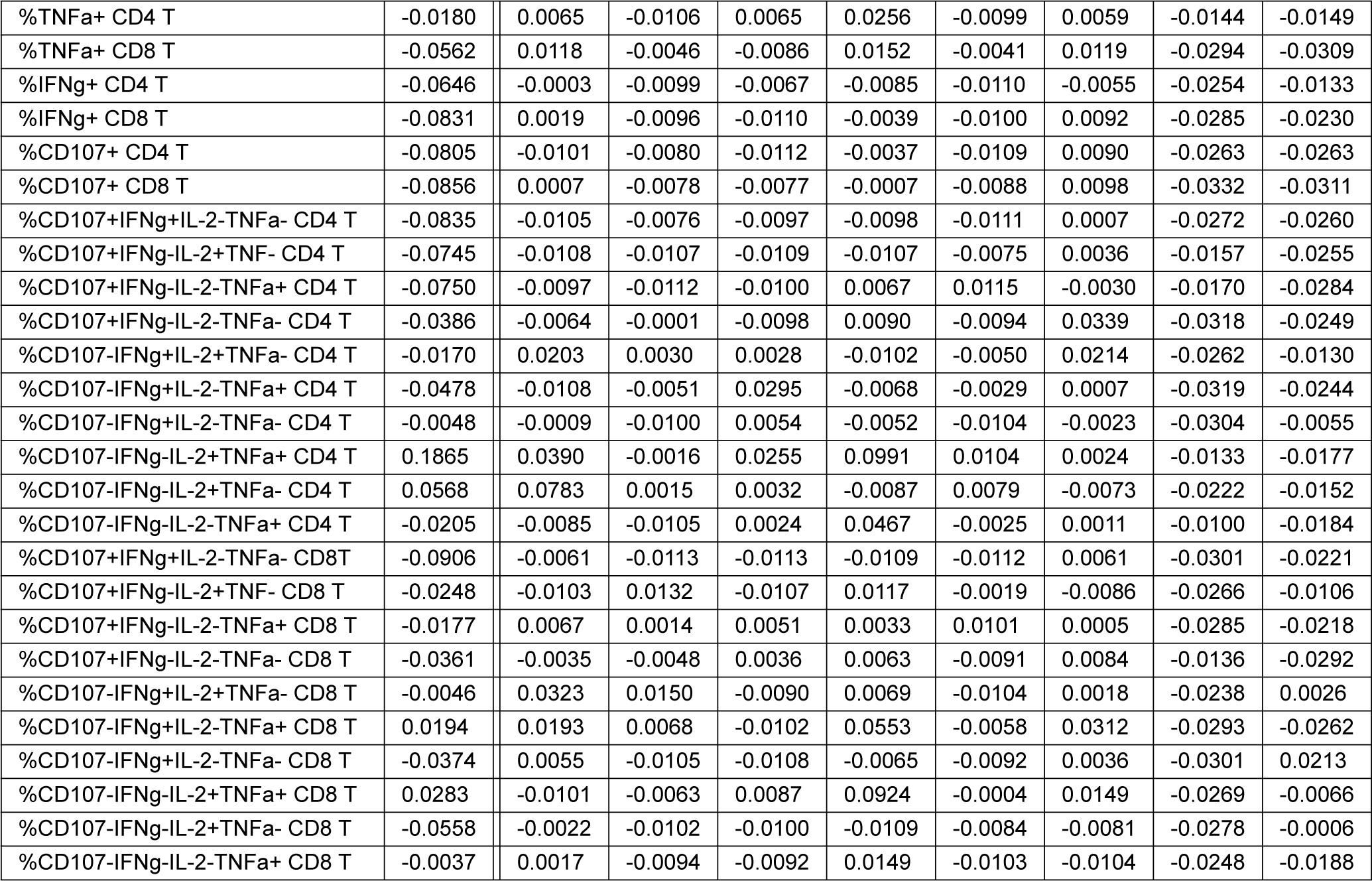
Adjusted R^2^ scores and differences in adjusted R^2^ for LOCO analysis for the model that contains total reservoir frequency. The first column contains all immunophenotypes and is a target for the regression model. The second column contains adjusted R^2^ of the linear regression model with features such as total reservoir frequency, age, biological sex, race, years of treatment, CD4 nadir, recent CD4 count, and years of HIV before treatment (=NA, *<* 1, *≥* 1). The next seven columns show differences in adjusted R^2^ score (Δ*R*^2^) after removing a specific feature. Participants with missing years of ART values are excluded from this analysis. The missing value of the CD4 nadir for one participant is imputed.

**Table S7:**
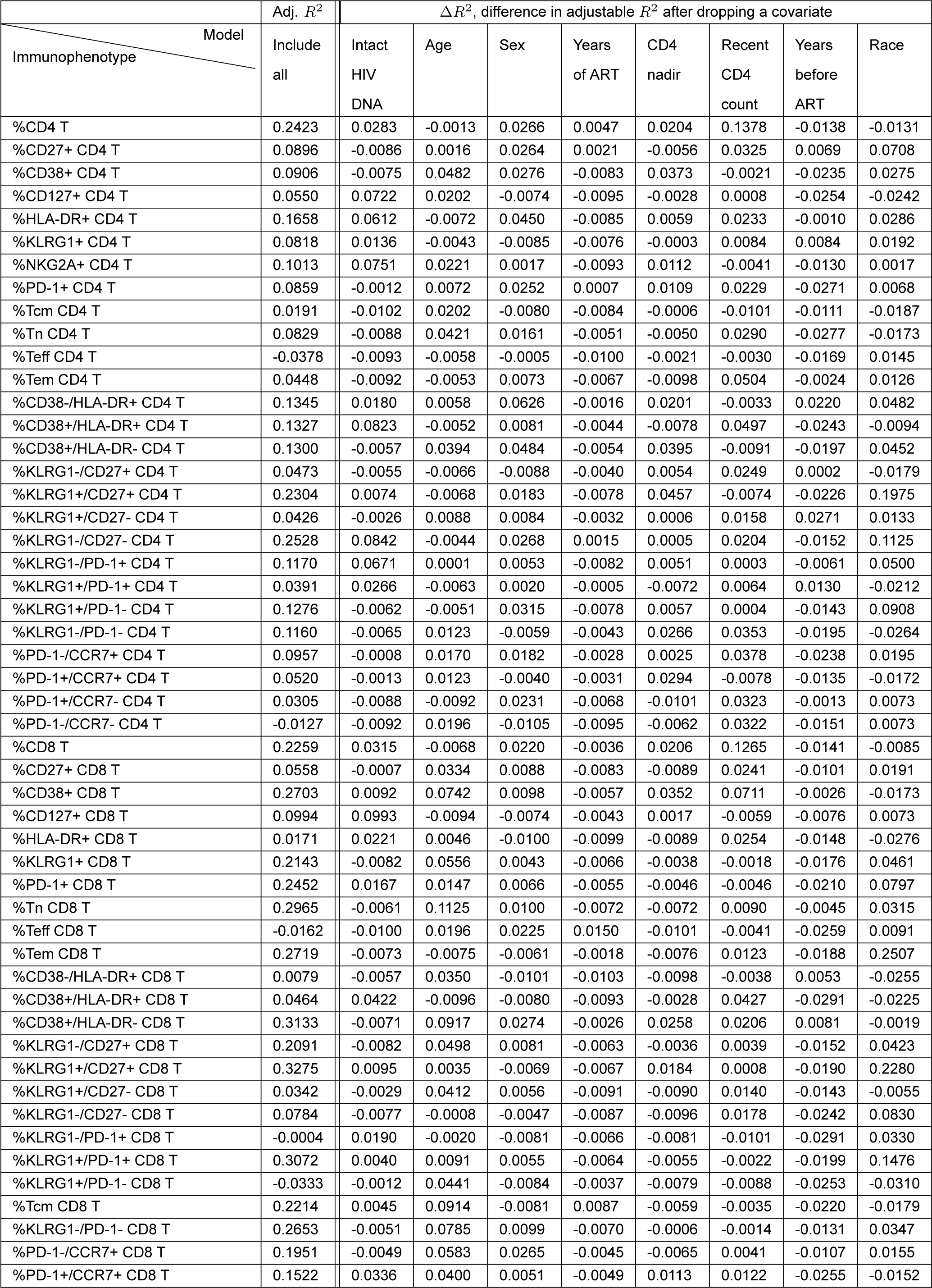
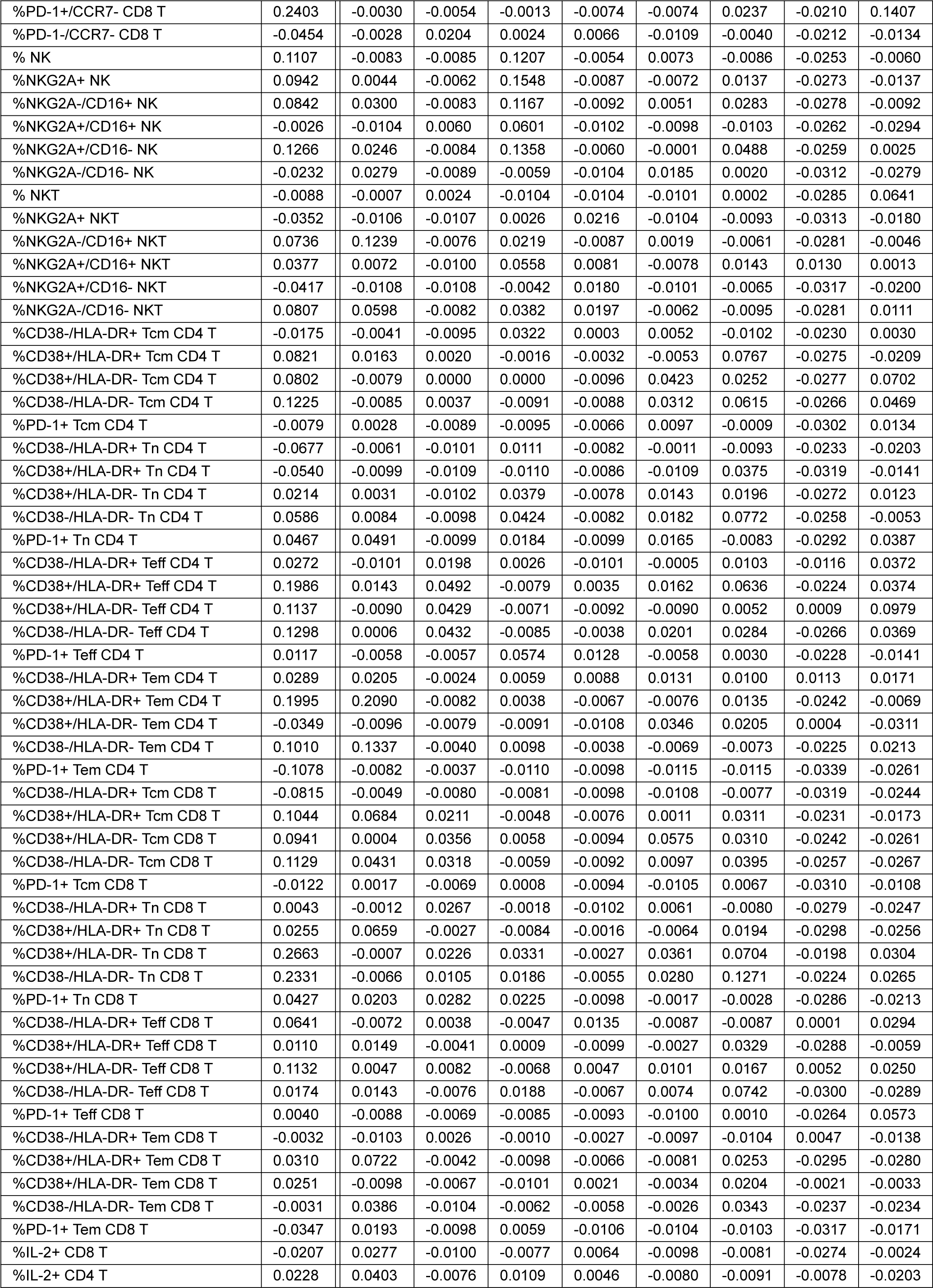
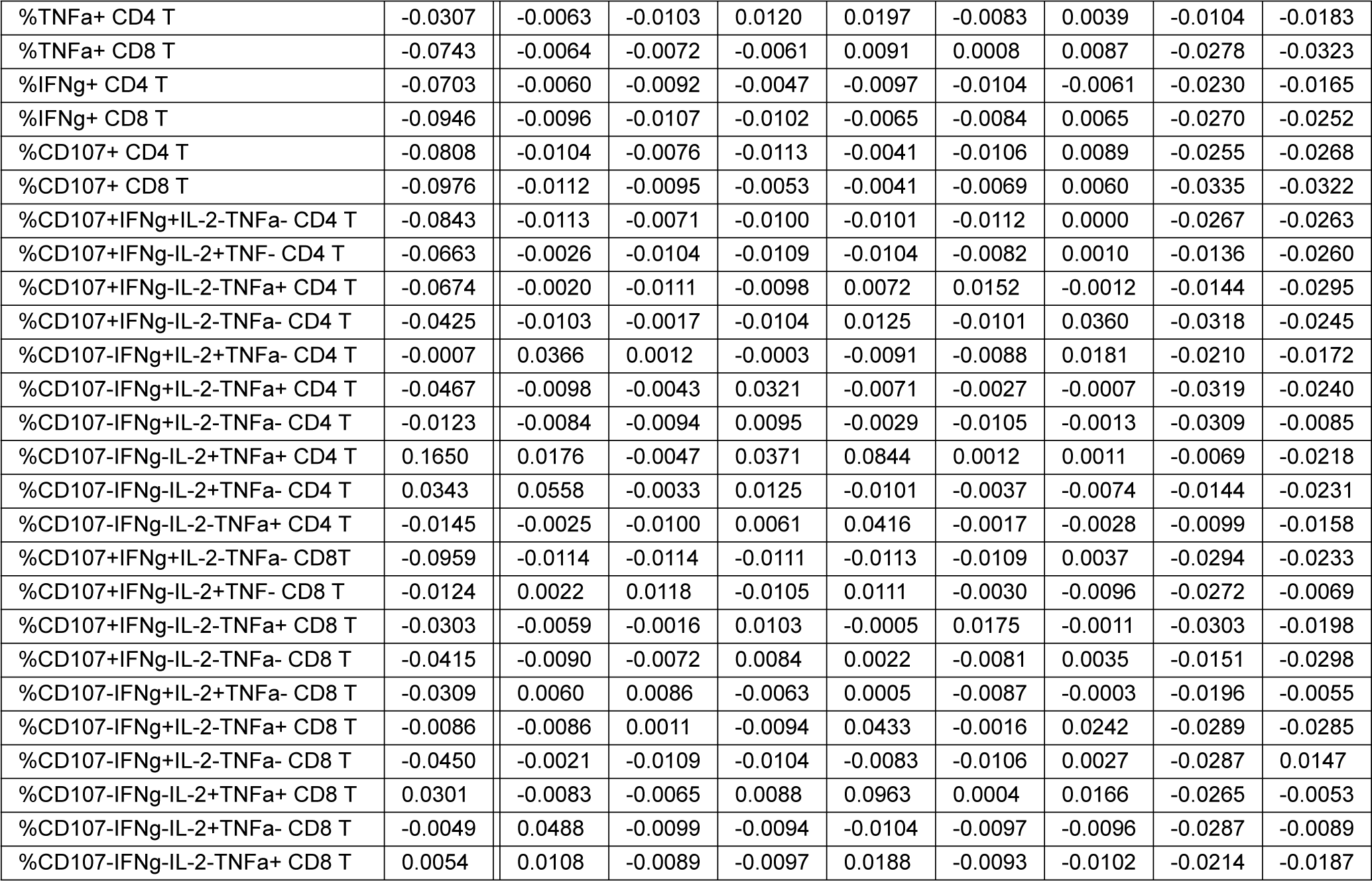
Adjusted R^2^ scores and differences in adjusted R^2^ for LOCO analysis for the model that contains intact reservoir frequency. The first column contains all immunophenotypes and is a target for the regression model. The second column contains adjusted R^2^ of the linear regression model with features such as intact reservoir frequency, age, biological sex, race, years of treatment, CD4 nadir, recent CD4 count, and years of HIV before treatment (=NA, *<* 1, *≥* 1). The next seven columns show differences in adjusted R^2^ score (Δ*R*^2^) after removing a specific feature. Participants with missing years of ART values are excluded from this analysis. The missing value of the CD4 nadir for one participant is imputed.

**Table S8:**
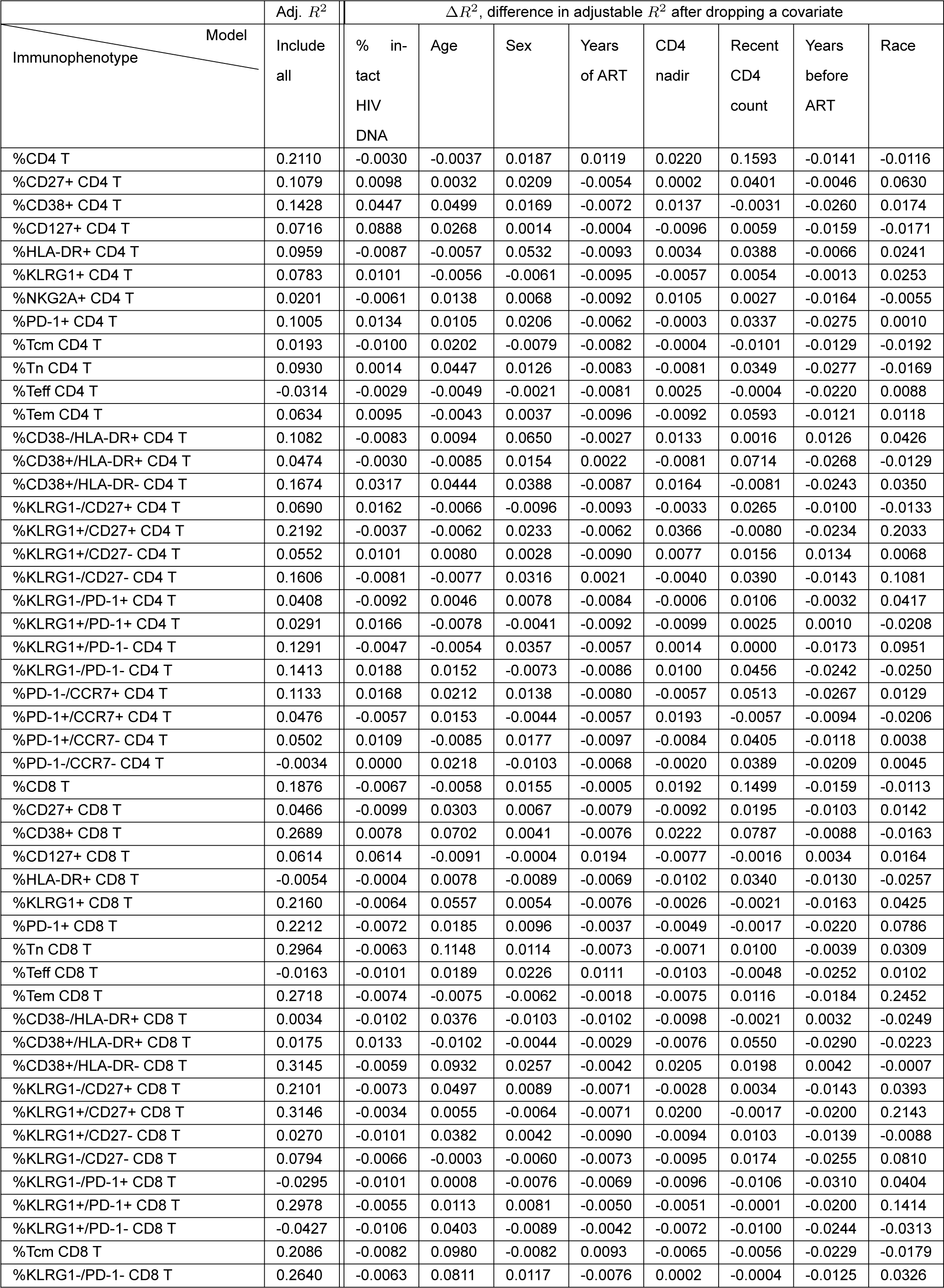
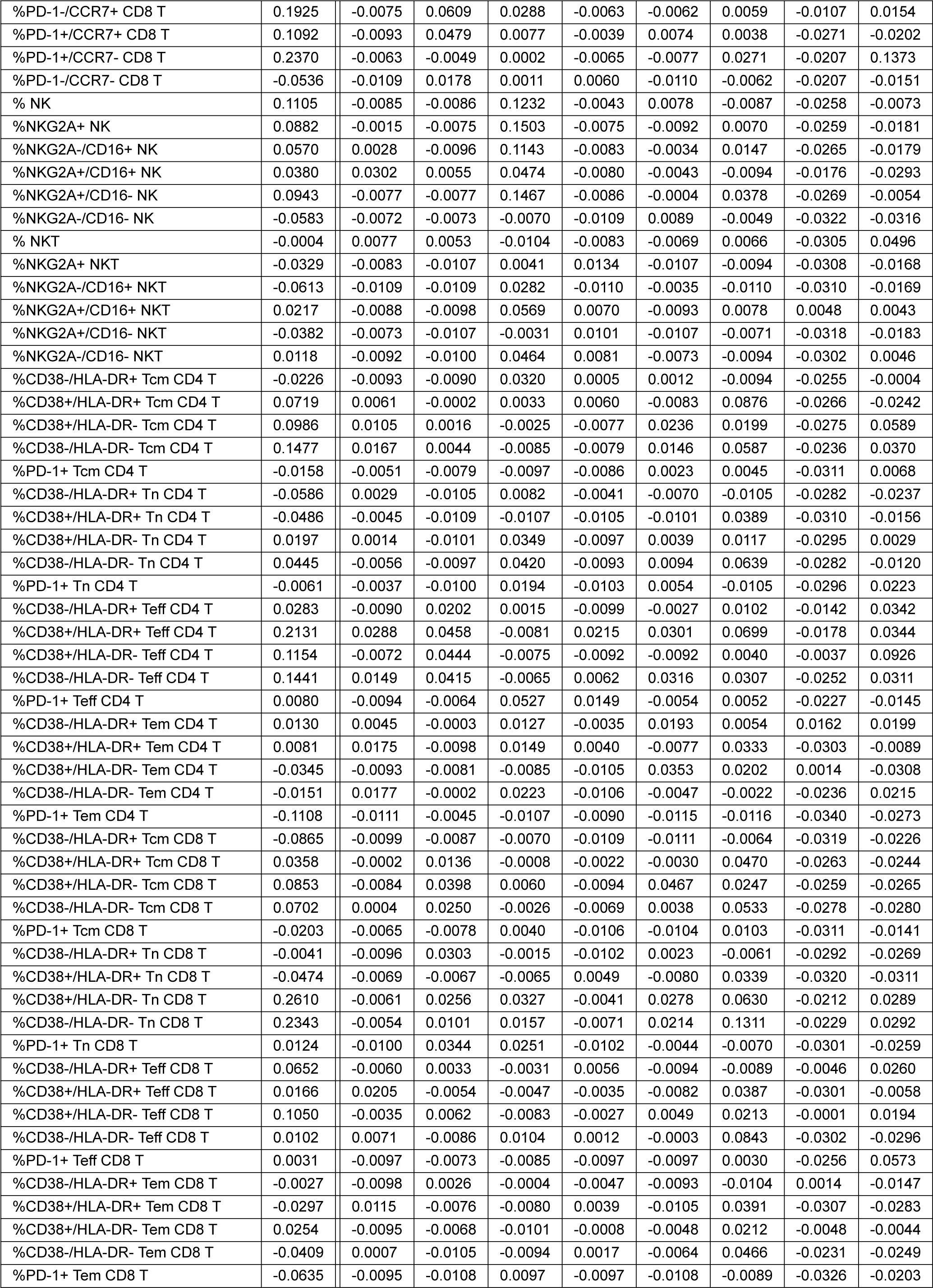
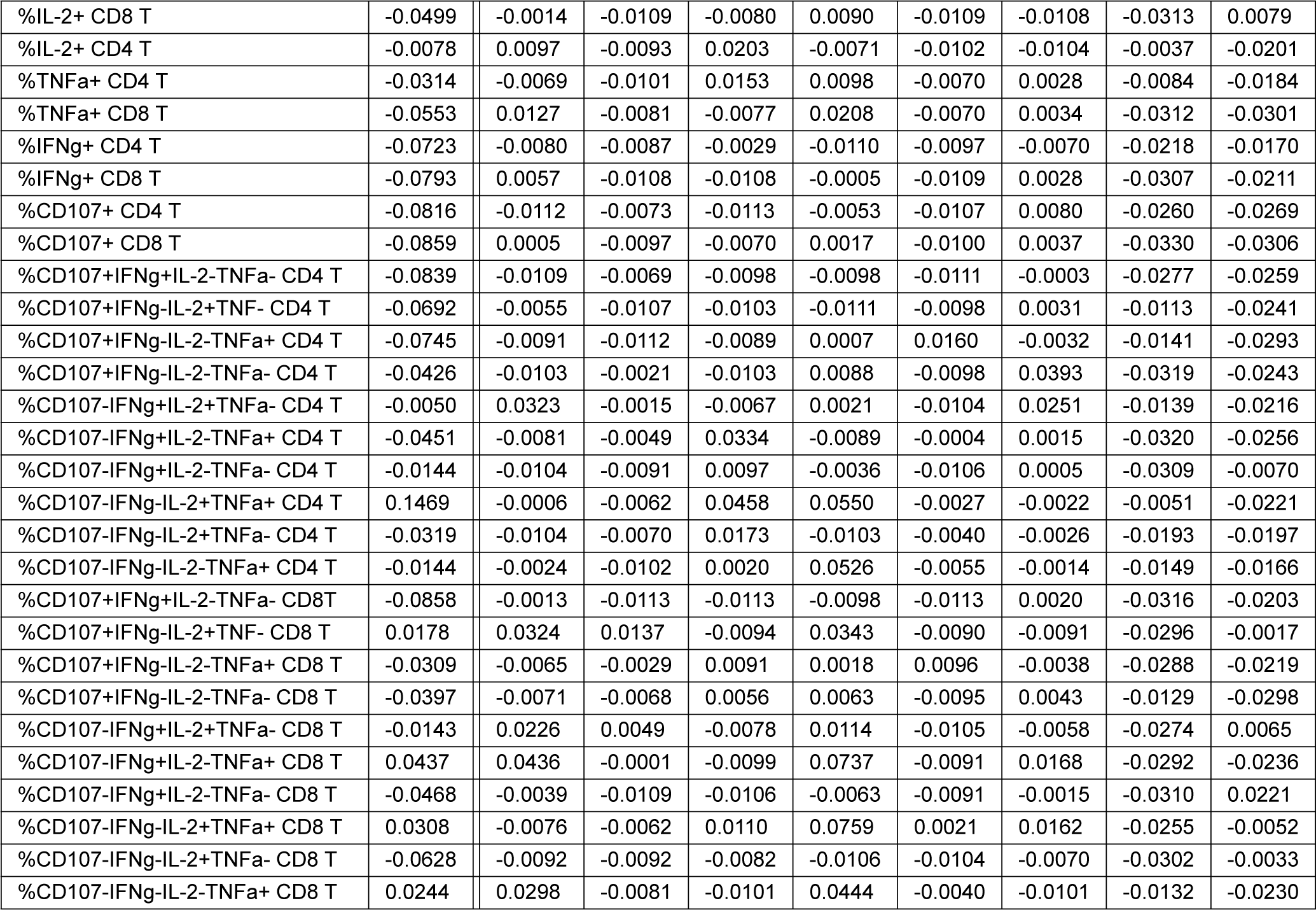
Adjusted R^2^ scores and differences in adjusted R^2^ for LOCO analysis for the model that contains percent intact. The first column contains all immunophenotypes and is a target for the regression model. The second column contains adjusted R^2^ of the linear regression model with features such as percent intact, age, biological sex, race, years of treatment, CD4 nadir, recent CD4 count, and years of HIV before treatment (=NA, *<* 1, *≥* 1). The next seven columns show differences in adjusted R^2^ score (Δ*R*^2^) after removing a specific feature. Participants with missing years of ART values are excluded from this analysis. The missing value of the CD4 nadir for one participant is imputed.

**Table S9:** Host features classify PWH with respect to HIV reservoir characteristics. The frequency of each of the 144 variables was analyzed for performance in classifying each PWH within the cohort as having reservoir frequency above or below the median using a receiver operating characteristics (ROC) curve. The area under the curve (AUC) is reported. The table includes variables that had an AUC value larger than 0.6 for one of the reservoir frequency characteristics. We highlight variables that had an AUC value higher than 0.65 in **bold**. Variables are ranked by AUC for the total reservoir frequency. For years of ART, AUC values are computed based on 108 PWH, excluding participants with missing years of ART values. For CD4 Nadir, AUC values are computed based on 114 PWH, excluding one participant] with the missing CD4 Nadir value.

| Variable | Bin Total HIV DNA | Bin Intact HIV DNA | Bin % intact HIV DNA | Variable | Bin Total HIV DNA | Bin Intact HIV DNA | Bin % intact HIV DNA |
| --- | --- | --- | --- | --- | --- | --- | --- |
| %NKG2A+ CD4 T | **0.6953** | 0.6084 | 0.5416 | %PD-1-/CCR7+ CD8 T | 0.6207 | 0.6134 | 0.5115 |
| %PD-1+ Tn CD4 T | **0.6827** | 0.5749 | 0.5313 | %CD38+ CD8 T | 0.6142 | 0.5425 | 0.6222 |
| %CD38+/HLA-DR- CD8 T | **0.6777** | 0.5797 | 0.6084 | %CD38-/HLA-DR- Teff CD4 T | 0.6122 | 0.5587 | 0.5017 |
| %Tn CD4 T | **0.6763** | 0.5933 | 0.5665 | %CD38-/HLA-DR+ Tn CD8 T | 0.6122 | 0.5420 | 0.5933 |
| %KLRG1-/PD-1- CD4 T | **0.6745** | 0.5789 | 0.6224 | %KLRG1-/CD27+ CD8 T | 0.6096 | 0.6101 | 0.5265 |
| %Tcm CD8 T | **0.6729** | 0.6205 | 0.5541 | %CD38+/HLA-DR- Tn CD4 T | 0.6078 | 0.5050 | 0.5531 |
| %CD38+/HLA-DR- CD4 T | **0.6700** | 0.5507 | 0.5889 | %CD107-IFNγ+IL-2-TNFα- CD8 T | 0.6065 | 0.5712 | 0.5133 |
| %CD8 T | **0.6629** | 0.6547 | 0.5484 | %CD38+/HLA-DR+ Tem CD4 T | 0.6057 | 0.5829 | 0.5941 |
| Age | **0.6627** | 0.6264 | 0.5476 | %CD38+/HLA-DR- Tcm CD8 T | 0.6039 | 0.5150 | 0.5750 |
| %PD-1+/CCR7+ CD8 T | **0.6608** | 0.5907 | 0.5328 | %CD38-/HLA-DR+ Tcm CD8 T | 0.6030 | 0.6053 | 0.5015 |
| %PD-1+ CD8 T | **0.6562** | 0.6207 | 0.5070 | %KLRG1-/PD-1+ CD4 T | 0.6019 | 0.5346 | 0.5290 |
| %PD-1+ Tn CD8 T | **0.6553** | 0.5880 | 0.5597 | %NKG2A-/CD16+ NK | 0.6012 | 0.5257 | 0.5848 |
| %Tn CD8 T | **0.6550** | 0.6485 | 0.5138 | %KLRG1+ CD8 T | 0.5954 | 0.6021 | 0.5251 |
| %PD-1-/CCR7+ CD4 T | **0.6537** | 0.5473 | 0.6178 | %KLRG1-/CD27+ CD4 T | 0.5945 | 0.5322 | 0.6205 |
| %PD-1+/CCR7+ CD4 T | 0.6475 | 0.5656 | 0.5740 | %CD27+ CD4 T | 0.5776 | 0.5018 | 0.6021 |
| %Tcm CD4 T | 0.6452 | 0.5998 | 0.5104 | %KLRG1+/CD27- CD4 T | 0.5723 | 0.5017 | 0.6181 |
| %KLRG1-/PD-1- CD8 T | 0.6440 | 0.6308 | 0.5234 | %PD-1+ Tcm CD4 T | 0.5705 | 0.5005 | 0.6158 |
| %PD-1+ CD4 T | 0.6434 | 0.5286 | 0.6295 | %PD-1+ Tem CD4 T | 0.5688 | 0.6124 | 0.5489 |
| %CD4 T | 0.6429 | 0.6338 | 0.5322 | %KLRG1+ CD4 T | 0.5649 | 0.5243 | 0.6236 |
| %CD107-IFNγ+IL-2-TNFα- CD4 T | 0.6420 | 0.5670 | 0.5185 | Recent CD4 count | 0.5620 | 0.6320 | 0.6104 |
| %CD38+ CD4 T | 0.6414 | 0.5299 | 0.5932 | %KLRG1+/PD-1+ CD4 T | 0.5602 | 0.5100 | 0.6450 |
| %CD127+ CD4 T | 0.6367 | **0.7051** | 0.6323 | %Tem CD4 T | 0.5576 | 0.5230 | 0.6145 |
| %CD38+/HLA-DR- Tn CD8 T | 0.6364 | 0.5558 | 0.5724 | %IL-2+ CD4 T | 0.5416 | 0.6022 | 0.5687 |
| %PD-1-/CCR7- CD4 T | 0.6340 | 0.6010 | 0.5473 | %CD107-IFNγ+IL-2+TNFα- CD4 T | 0.5342 | 0.6308 | 0.6248 |
| %KLRG1+/PD-1+ CD8 T | 0.6335 | 0.6098 | 0.5020 | %CD38+/HLA-DR- Teff CD8 T | 0.5306 | 0.5613 | 0.6276 |
| CD4 Nadir | 0.6303 | 0.6110 | 0.5563 | Years before ART < 1 | 0.5299 | 0.5222 | 0.6092 |
| Years of ART | 0.6296 | 0.5069 | **0.7171** | %CD127+ CD8 T | 0.5195 | 0.6051 | 0.6048 |
| %CD38-/HLA-DR- Tem CD4 T | 0.6249 | 0.5513 | 0.5867 | %CD107-IFNγ-IL-2+TNFα+ CD4 T | 0.5003 | 0.6000 | 0.6493 |
| %KLRG1+/CD27+ CD8 T | 0.6224 | 0.6473 | 0.5153 |  |  |  |  |

**Figure S8:**
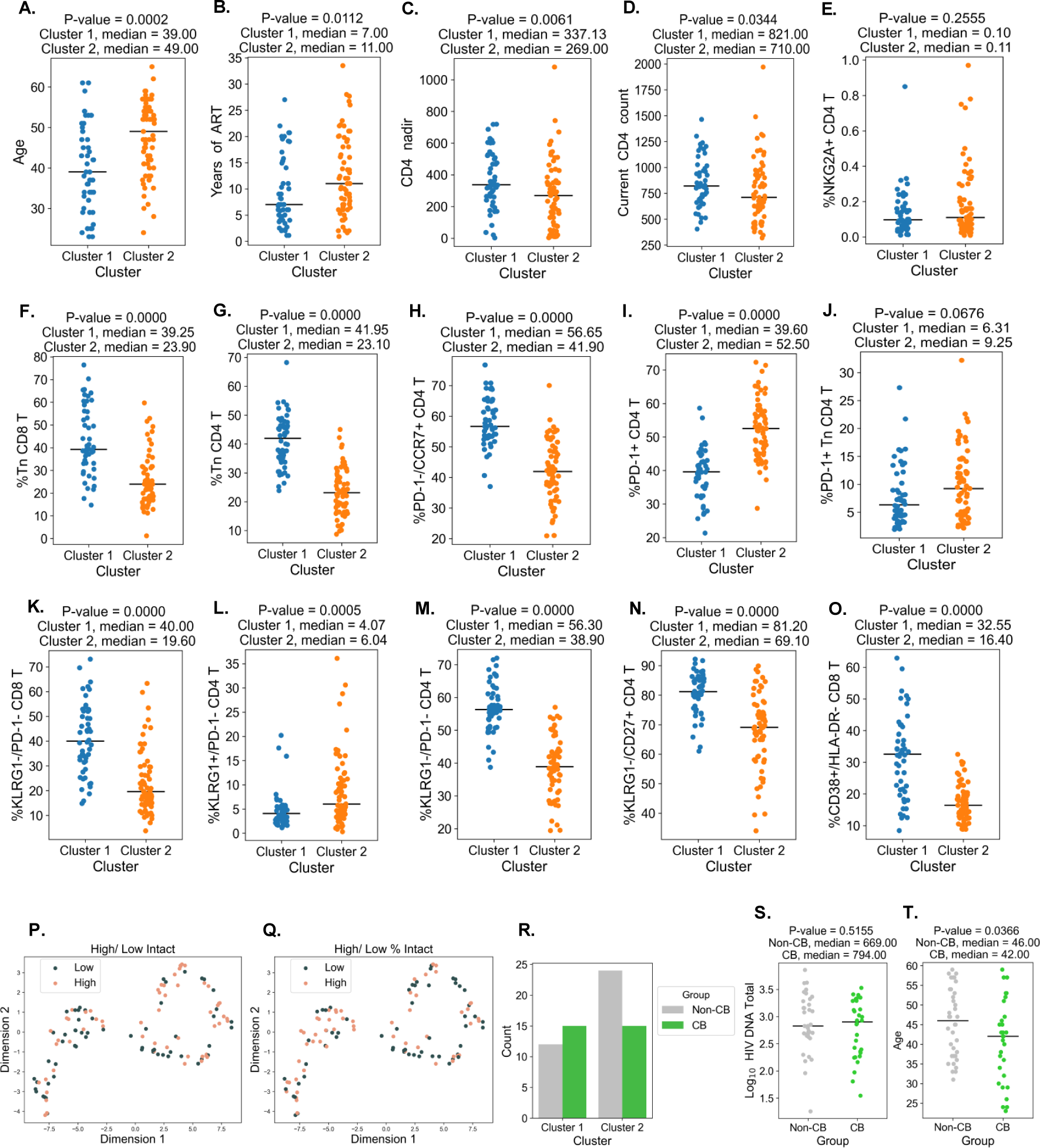
Dimension reduction supplemental figures. **A.** Participant age is shown within each cluster. **B.** Participant years of ART are shown within each cluster. **C.** Participant CD4 nadir is shown within each cluster. **D.** Participant CD4 count is shown within each cluster. **E-O**. Participant immune features of interest are shown, where plots of immune features with similar names are placed nearby. **P.** Clusters with data points color-coded by intact reservoir frequency (high=pink, low=gray). **Q.** Clusters with data points color-coded by percent intact reservoir frequency (high=pink, low=gray). **R.** Relative proportions of cannabis (CB) users and non-users (non-CB) are shown for each cluster. **S.** Total reservoir frequencies (per million CD4 T cells) for non-users and CB users are shown. **T.** The ages of study participants for non-users and CB users are shown.

**Figure S9:**
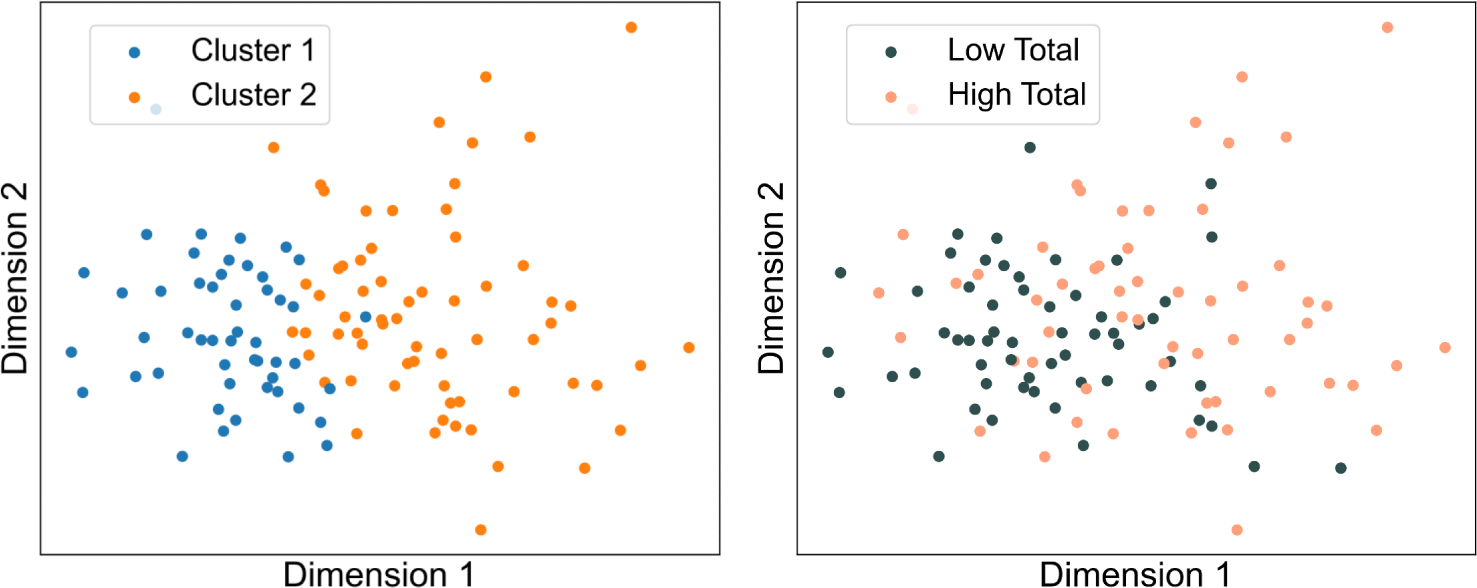
**Left:** PCA plot with data points color-coded by the membership of clusters identified in **Figure 4**. **Right:** PCA plot with data points color-coded by total reservoir frequency (high=pink, low=gray).

**Figure S10:**
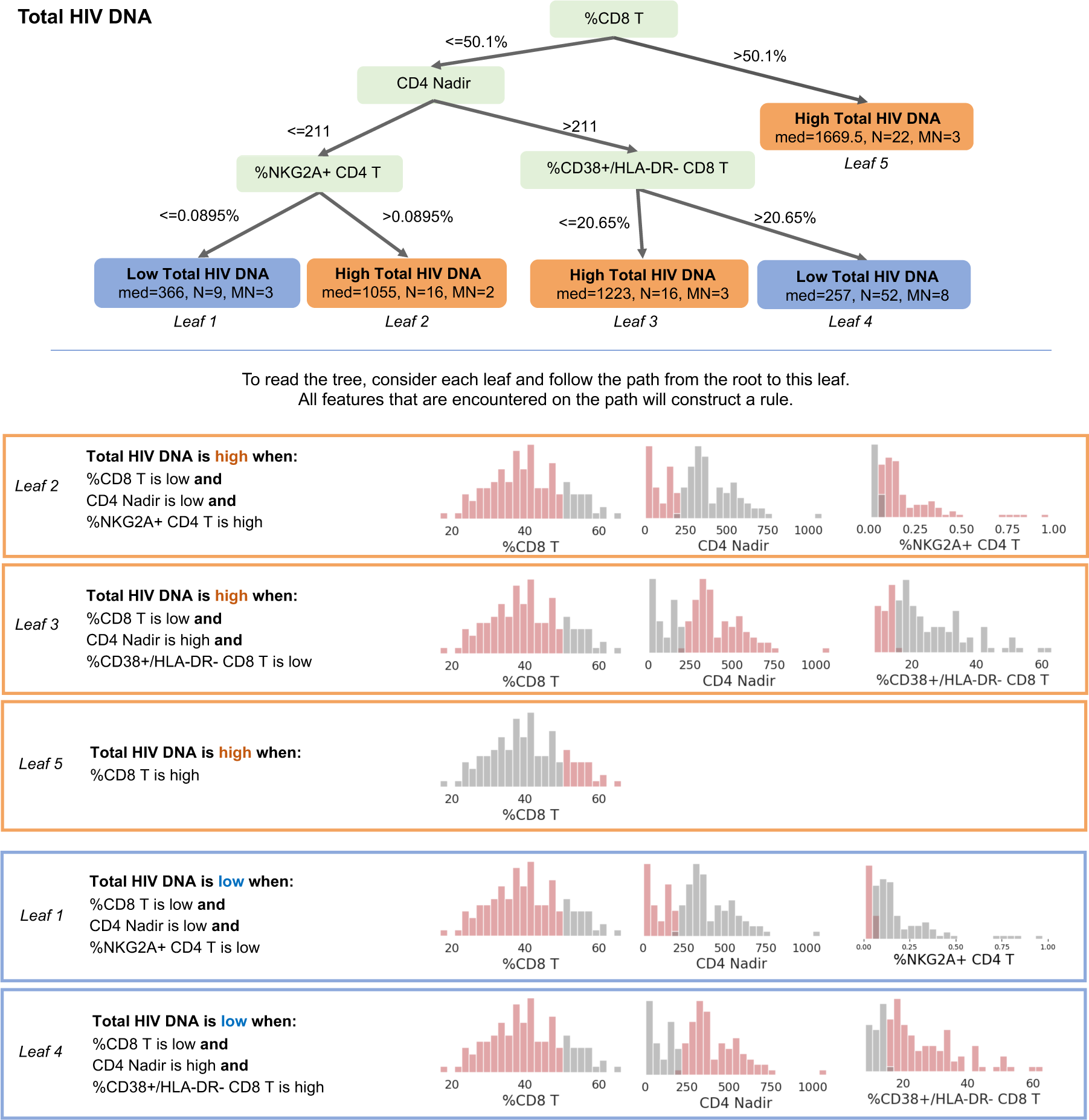
The total reservoir frequency visualization tree from Figure 6 is explained with a set of rules. For every leaf, the path that leads to this leaf is described. The histogram of the variable and split value for every node used in the tree is shown.

**Figure S11:**
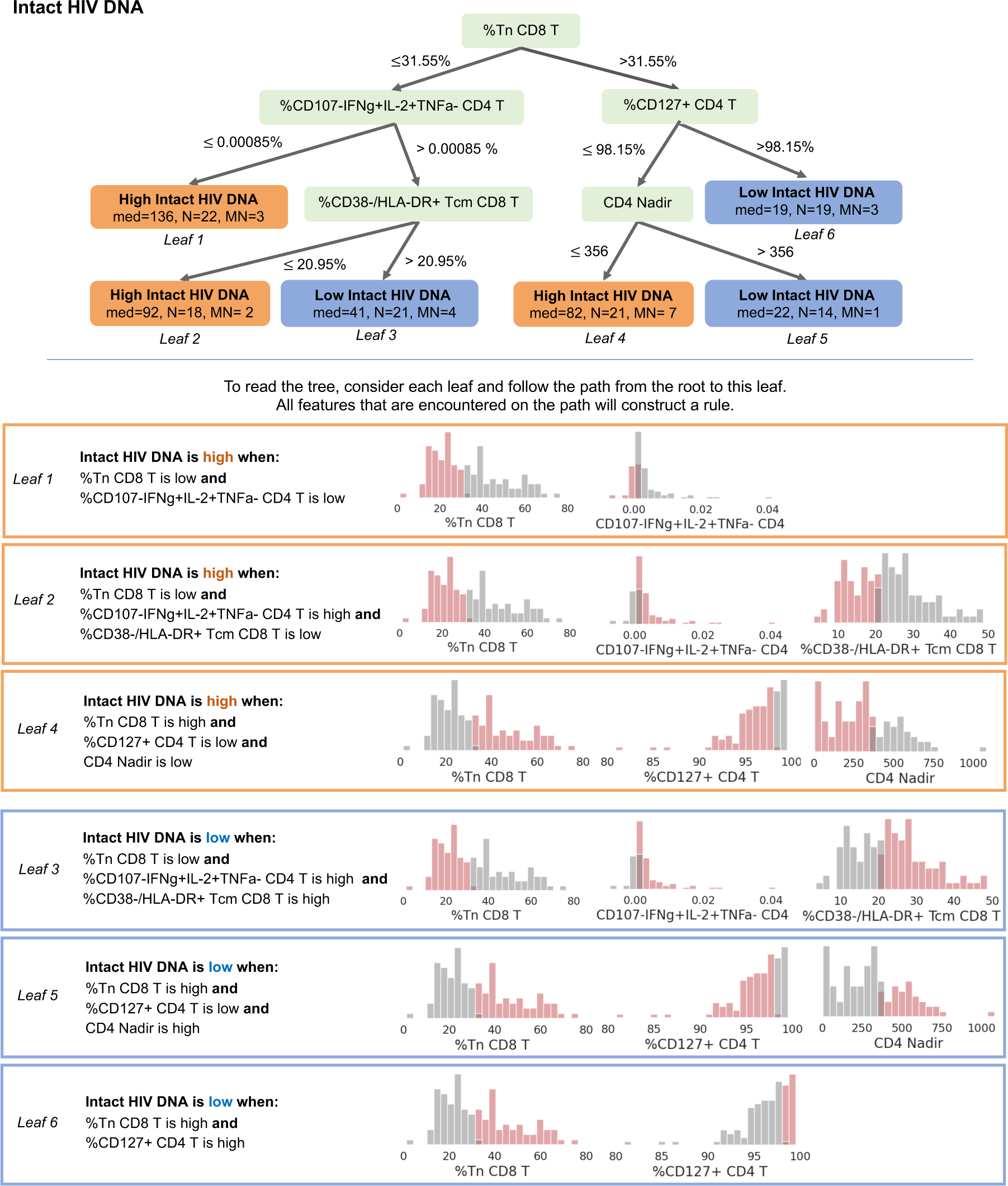
The intact reservoir frequency visualization tree from Figure 6 is explained with a set of rules. For every leaf, the path that leads to this leaf is described. The histogram of the variable and split value for every node used in the tree is shown.

**Figure S12:**
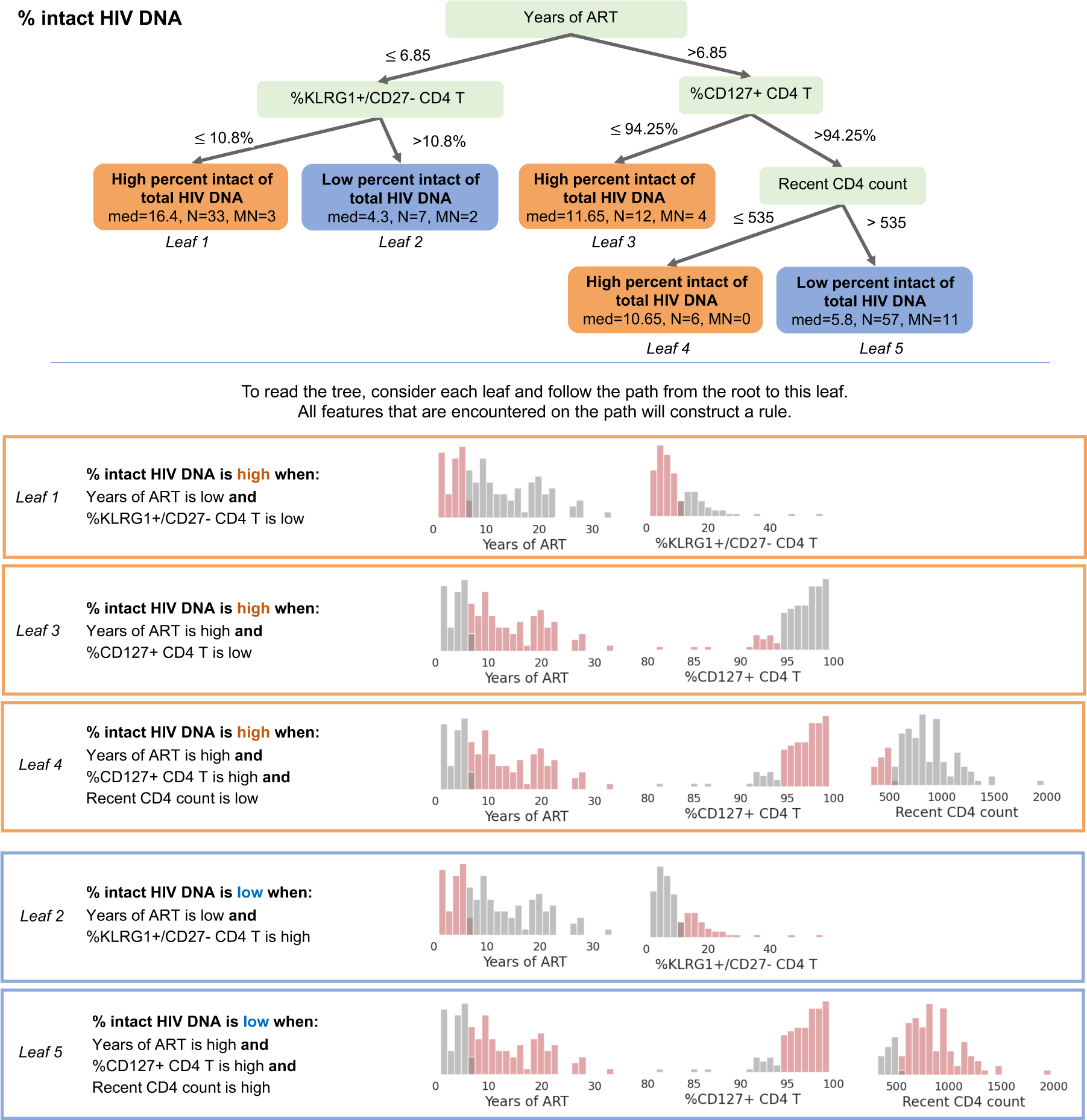
The percent intact visualization tree from Figure 6 is explained with a set of rules. For every leaf, the path that leads to this leaf is described. The histogram of the variable and split value for every node used in the tree is shown.

**Figure S13:**
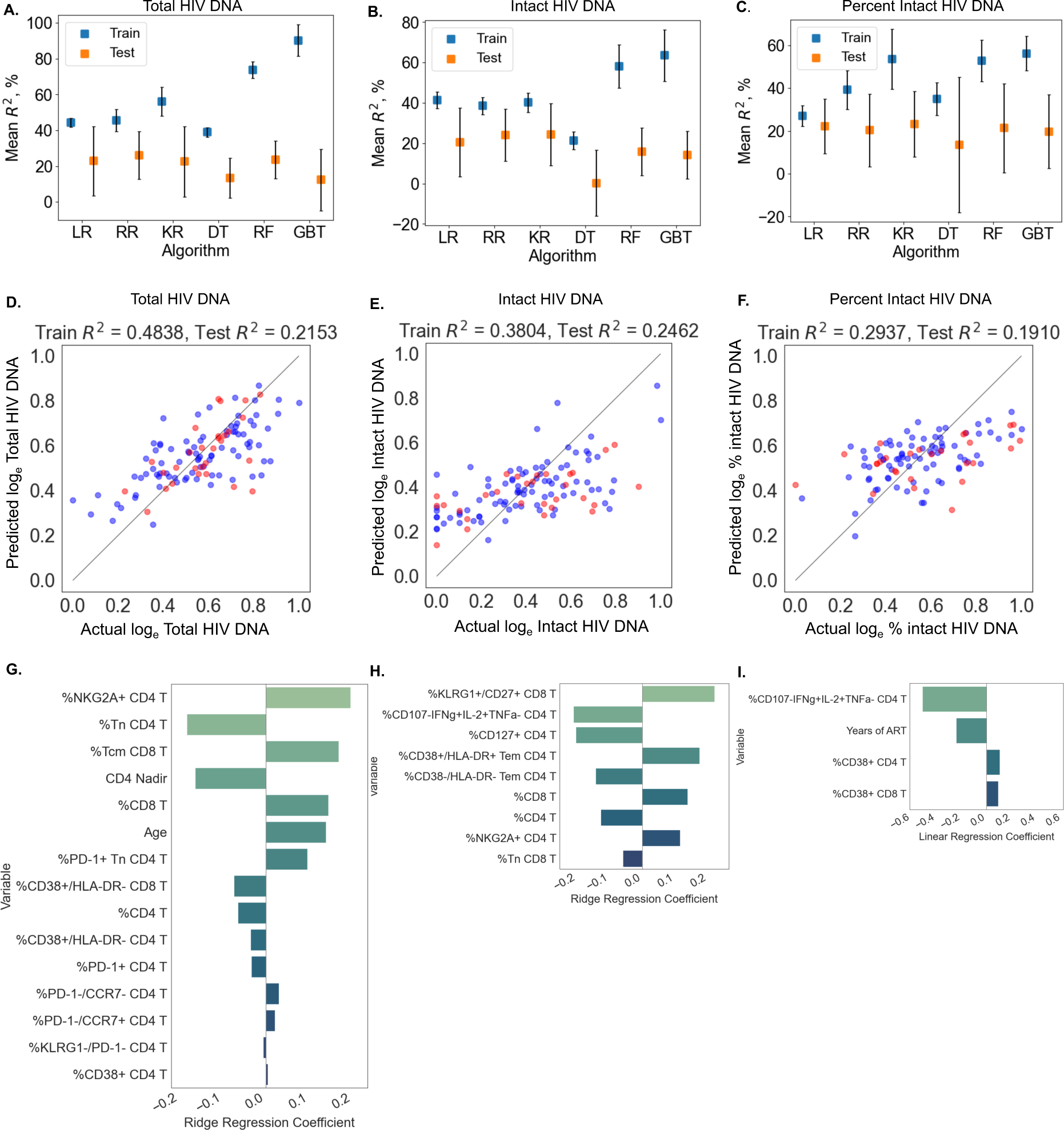
Using machine learning to predict reservoir frequency. Average training and test R^2^ scores over different training and test data splits for Linear Regression (LR), Ridge Regression (RR), Kernel Regression with RBF kernel (KR), Decision Tree Regressor (DT), Random Forest (RF), and Gradient Boosted Trees (GBT) models are shown for predicting total reservoir frequency (**A**), intact reservoir frequency (**B**), and percentage intact (**C**). For one split of training and test sets, ridge regression performance is shown for total reservoir frequency (**D**), intact reservoir frequency (**E**), and linear regression for percentage intact (**F**). On the x-axis, the actual values of HIV reservoir characteristics are shown, while on the y-axis the outputs of models are shown for training data (blue dots) and test data (red dots). For the same training and test split, ridge regression model feature coefficients are visualized for total reservoir (**G**), intact reservoir (**H**), and linear regression model feature coefficients are visualized for percent intact (**I**). On the y-axis we show variables used by the model, while on the x-axis coefficient values for linear models based on these variables. For percent intact models we exclude participants with missing values of years of ART. For total and intact reservoir frequency, missing values of years of ART were imputed. The missing value of the CD4 nadir for one participant was imputed as well using the MICE algorithm.

## Algorithm 1 Training procedure for classification (regression)

**Input:** Outcome, algorithm, set of hyperparameters from Table S10

**Output:** Mean and standard deviation of train and test accuracies (R^2^ scores)

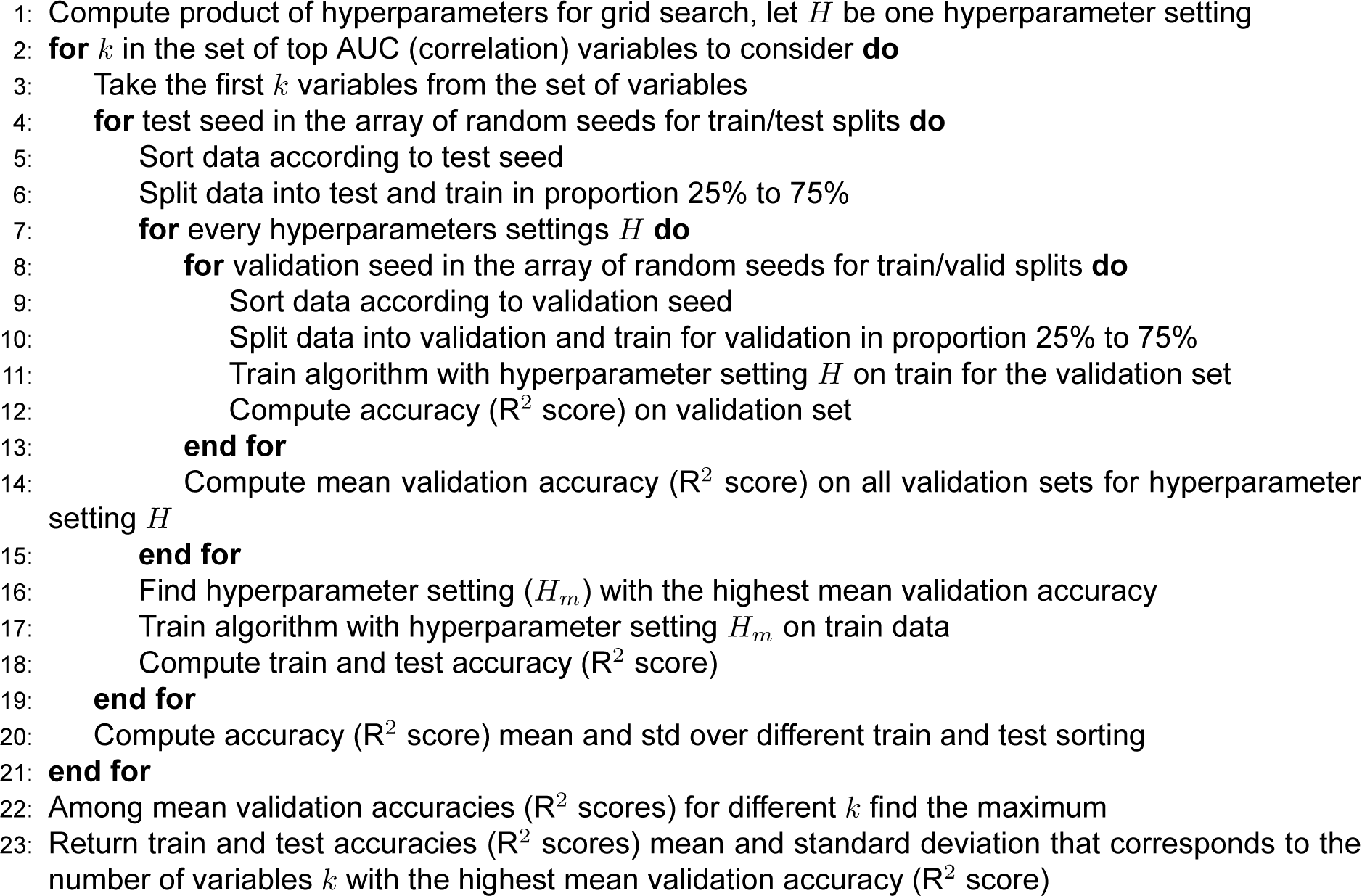

**Table S10:** Ranges of hyperparameters values that we used to perform grid search for classification and regression.

|  |  |
| --- | --- |
| **General** |  |
| Random seeds for train/test splits | 0..10 |
| Random seeds for train/valid splits | 0..20 |
| Set of the top AUC variables to consider | 2..9,10,15,20,25,30,40,50 |
| Set of the top absolute value of correlation coefficient to consider | 2..9,10,15,20,25,30,40, all |
| **Logistic Regression** (classifier) |  |
| Penalty | L2 |
| Regularization parameter | 0.00001, 0.0001, 0.001, 0.01, 0.1, 1, 10, 100, 1000, 10000 |
| **SVM with RBF kernel** (classifier) |  |
| Regularization parameter | 0.00001, 0.0001, 0.001, 0.01, 0.1, 1, 10, 100, 1000, 10000 |
| **Ridge Regression** (regressor) |  |
| Regularization parameter | 0.00001, 0.0001, 0.001, 0.01, 0.1, 1, 10, 100, 1000, 10000 |
| **Kernel Ridge Regression** (regressor) |  |
| Regularization parameter | 0.00001, 0.0001, 0.001, 0.01, 0.1, 1, 10, 100, 1000, 10000 |
| Kernel parameter | 0.00001, 0.0001, 0.001, 0.01, 0.1, 1, 10, 100, 1000, 10000 |
| **CART** (classifier) and **Decition Tree** (regressor) |  |
| Maximum depth | 2..7, 10 |
| Minimum number of samples in the leaf | 4, 6, 8, 10 |
| **Random Forest and Gradient Boosted Trees** (classifier and regressor) |  |
| Maximum depth | 2..7, 10 |
| Minimum number of samples in the leaf | 4, 6, 8, 10 |
| Maximum number of estimators | 2, 5, 10, 20, 50, 100, 200 |

